# Expanding growers’ choice of disease management options can promote suboptimal social outcomes

**DOI:** 10.1101/2022.09.05.506581

**Authors:** Rachel E. Murray-Watson, Nik J. Cunniffe

**Affiliations:** Department of Plant Sciences, University of Cambridge, Cambridge, United Kingdom

**Keywords:** behavioural model, epidemiological model, Pareto optimisation, Gini coefficient, TYLCV

## Abstract

Previous models of growers’ decision-making during epidemics have unrealistically limited disease management choices to just two options. Here, we expand previous game-theoretic models of grower decision-making to include three control options: crop that is either tolerant, resistant, or susceptible to disease. Using Tomato Yellow Leaf Curl Virus (TYLCV) as a case study, we investigate how growers can be incentivised to use different control options to achieve socially-optimal outcomes. To do this, we consider the efforts of a “social planner” who moderates the price of crops. We find that subsidising tolerant crop costs the social planner more in subsidies, as its use encourages selfishness and widespread adoption. Subsidising resistant crop, however, provides widespread benefits by reducing the prevalence of disease across the community of growers, including those that do not control, reducing the number of subsidies required from the social planner. We then use Gini coefficients to measure equitability of each subsidisation scheme. This study highlights how grower behaviour can be altered using crop subsidies to promote socially-optimal outcomes during epidemics.

## 2 Introduction

Those tasked with managing epidemics typically have limited resources which they must distribute efficiently. This often incurs a trade-off involving multiple parties with conflicting objectives, and how these trade-offs are managed will depend on the relative priority of each objective. A well-known example of such a trade-off for disease management occurs during vaccination campaigns (e.g. Gavish & Katriel (2022), Yousefpour et al. (2020)). Members of the public often wish to minimise their infection risk, whilst governments may want to optimise the allocation of vaccines to protect the most vulnerable (Gavish & Katriel (2022)). Similarly, in the context of plant epidemics, growers faced with crop disease will want to maximise profits, but with this comes potential environmental damage (Zaffaroni & Bevacqua (2022)) or reduced profitability when there is no risk of infection (Vyska et al. (2016)). The control mechanism adopted by growers may also have negative consequences for others; Grogan & Goodhue (2012) found that when growers sprayed pesticides within their non-citrus fields, the pesticides killed parasitic wasps that are the natural enemy of pests in neighbouring citrus fields. This intensified the reliance of citrus growers on pesticides, increasing their expenditure and environmental impact.

In economics, a social planner is a decision-maker whose goal is to balance the trade-offs that result from a policy action, aiming to maximise the welfare across all parties (Sugden (2013a)). Originally conceptualised for welfare economics (Sugden (2013b)), the social planner’s problem has been used in a range of settings, such as infectious disease management (Gersovitz & Hammer (2005), Toxvaerd (2019), Giannitsarou et al. (2021), Toxvaerd & Rowthorn (2022)), wildlife conservation (Johannesen & Skonhoft (2005)) and supply chain management (Lee & Choi (2021)). A key aspect of the social planner is that they account for the externalities associated with certain actions. In economics, externalities are the consequences of an action that are felt by a third party not involved in the action (Gersovitz (2014)). These externalities are often ignored by individuals, who only care about their own outcome, but they are important when considering what is socially optimal.

The social planner’s problem can be solved using Pareto optimality. Under a Pareto-optimal solution, we cannot improve one party’s objective without ensuring a worse outcome for another party (Luc (2008)). These solutions are efficient, as they optimise for each outcome, and non-dominated, as, given the same constraints, they cannot be improved upon by any other solution. There may be many possible solutions for a given problem, which combine to give a Pareto front. Pareto fronts have been calculated to visualise the trade-offs involved in the recent coronavirus pandemic (Yousefpour et al. (2020)), to retrospectively evaluate responses to the 1918 influenza pandemic (Velde (2022)) and to better understand how to optimally allocate healthcare equipment during an epidemic (Donmez et al. (2022)). In plant disease modelling, they have been used to investigate the trade-offs between crop productivity and environmental impacts (Zaffaroni & Bevacqua (2022)). None of the solutions in the Pareto front is inherently better than any other, so the decision-maker must decide which option to use based on additional information.

Here we use Pareto fronts to study the social planner’s problem in the context of a plant disease epidemic. The individuals in a population of growers can choose between different crop types that vary in their losses in yield if they become infected, their susceptibility to infection and/or their infectivity. The behavioural component of the model is based on game theory, an economic tool used to study the strategic decisionmaking undertaken by individuals when their choice of action strongly depends on the actions of others (Morris (2012)). Previous work that incorporated human behaviour into plant epidemic models has only allowed growers two options, “control” or “not control” (Milne et al. (2016), McQuaid et al. (2017a), Saikai et al. (2021), Milne et al. (2020), Bate et al. (2021), Murray-Watson et al. (2022) and Murray-Watson & Cunniffe (2022)), but such a narrow choice is unlikely, and in practice, growers will instead choose from a range of control options, each with different characteristics.

We adapt the game-theoretic models described in Milne et al. (2016) and subsequent literature to allow growers an expanded strategy set. We extend the model to go beyond the previous choice between “control” and “not control” to allow growers to choose between three crop types: two control options (disease-resistant or disease-tolerant crop) or unimproved crop. Growers’ decisions on which crop type to use will depend on their estimation of the profit they expect to earn over the next growing season if they were to use each crop type, which in turn depends on the prevalence of infection and the proportion of other growers using each control strategy. The models are based on “strategic adaptive” expectations (Fenichel & Wang (2013)), where growers balance their knowledge of the current state of the system with their experience from the previous season. In particular, growers compare their profit from the previous season with the expected profit over the next season for each of the strategies that they did not use (the “strategy vs. alternative” models described in Murray-Watson et al. (2022) and Murray-Watson & Cunniffe (2022)). This allows growers to balance sources of information, acting as “reflexive producers” (Kaup (2008)).

Resistance and tolerance are the two main molecular mechanisms of plant disease defence. Disease-resistant plants can limit pathogen replication, whilst tolerant hosts are not damaged as severely by the pathogen (Pagán & García-Arenal (2018)). Both traits lie on a spectrum, and in practice plants are more likely to have only partial (“quantitative”) disease resistance or tolerance (Marchant et al. (2020), French et al. (2016), Corwin & Kliebenstein (2017), Taylor & Cunniffe (2022)). Previous work has shown that tolerant and resistant crops caused contrasting outcomes for non-controllers (Murray-Watson & Cunniffe (2022)). Planting tolerant crop was often more beneficial for controllers than resistant crop, as it limited the yield loss in infected fields. However, their reduced symptom expression additionally meant that tolerant crops could not be rogued as effectively, leading to a build-up of inoculum and high infection levels. This lowered yields for all non-controllers. These effects directly incentivised use of the tolerant crop, as growers could expect to earn higher profits using tolerant crop than when they used the unimproved variety.

Conversely, as resistant crops are less susceptible to infection and limit pathogen replication once infected, the overall disease pressure on other fields is reduced when resistant crop is used. This increases the profits of non-controllers, as they can gain some of the benefits of resistant varieties without themselves paying the cost of the improved variety (i.e. can *free-ride* off controllers). The broader consequences of each crop type - the increase and decrease in infection pressure and the subsequent effect on profits of non-controllers - can therefore be considered *externalities* of each control decision. In this previous work, growers could not choose directly between resistant and tolerant crop, so the impact of these externalities on growers’ disease management practices when growers can choose between three crop types - unimproved, tolerant or resistant - is unknown.

As a case study, we study the effect of this three-way choice using *Tomato Yellow Leaf Curl Virus* (TYLCV). TYLCV is a global threat to tomato *(Lycopersicon esculentum*) production, and it has been detected in Australia, North America, Europe and East Asia (Ramos et al. (2019)). The virus is transmitted by *Bemisia tabaci* (Pan et al. (2012)) and infection causes leaf curling and chlorosis, stunting and up to 100% yield loss (Levy & Lapidot (2008), Dhaliwal et al. (2020)). Recently, tomato varieties that are either tolerant or resistant to TYLCV have been deployed (Lapidot et al. (1997), Riley & Srinivasan (2019)), helping to reduce yield loss.

We consider the perspective of a social planner who has the ability to subsidise these crop types. Historically, the European Union has subsidised the cultivation of processing tomato varieties (Sumner et al. (2001)); here, we study subsidies as a means of promoting particular epidemic outcomes. Subsidies are a means of rewarding/penalising a grower based on the external effects of their actions (i.e. internalising the externalities generated by using a particular practice) (Pretty et al. (2001)). They have a long history in agriculture, from regulating food production to promoting more environmentally-friendly practices such as hedgerow conservation in the UK (Stokstad (2020)) or reduced pesticide use in France (Aubert & Enjolras (2022)). They also have been used to encourage the use of particular crop varieties or control mechanisms (Carriere et al. (2020), Holden & Fisher (2015), Zhao et al. (2022), Okechukwu & Kumar (2016)). Here we consider two subsidies: one for resistant crop and one for tolerant crop, either of which is provided to the growers by a “social planner” or centralised body. As, in our model, growers choose between different crop types based on their expected profitability, subsidising each crop type incentivises its use. The planner has two objectives: to maximise the average profit across growers and to minimise their own spending on subsidies. We study the options available to the social planner using Pareto optimality.

The work, therefore, addresses three primary questions: 1) What are the longterm outcomes when growers have access to three different crop types? 2) How do the responses depend on the efficacy and cost of tolerant and resistant traits? 3) Which subsidisation strategy promotes socially-optimal solutions?

## 3 Methods

### 3.1 Epidemic model with three crop varieties

The following model extends Murray-Watson et al. (2022). It describes the spread of *Tomato Yellow Leaf Curl Virus* (TYLCV) through a population of fields, each cultivated by a single grower. The fields can be planted with one of three crop varieties: unimproved crop (*U*), crop that is tolerant to TYLCV (*T*) or crop that is resistant to TYLCV (R). We also track each field’s infection status; as we do not model within-field spread, fields can either be susceptible to infection (*S*), latently infected (E) or infectious (*I*). The parameterisation of this model is identical to that presented in Murray-Watson & Cunniffe (2022), with a summary of the parameter values presented in Tables 1 and 2. State variables are scaled to be a proportion of the total number of fields (i.e. *N* = *S_U_* + *E_U_* + *I_U_* + *S_T_* + *E_T_* + *I_T_* + *S_R_* + *E_R_* + *I_R_* = 1).

**Table 1:** Parameters related to resistant and tolerant varieties. The distinction between the two varieties lies in how disease is transmitted and what losses are incurred when a field is infectious. Here, *b* is the strategy of the grower and *b* ∈ {*U,T, R*}

| Parameter | Meaning | Value if<br>unimproved | Value if<br>resistant | Value if<br>tolerant |
| --- | --- | --- | --- | --- |
| $\delta_{\beta_b}$ | Relative susceptibility | 1 | 0.5 | 1 |
| $\delta_{\sigma_b}$ | Relative infectivity | 1 | 0.5 | 1 |
| $\delta_{Y_b}$ | Relative yield | 1 | 1 | 1 |
| $\delta_{L_b}$ | Relative losses due to<br>disease | 1 | 1 | 0.1 |
| $\frac{1}{\delta_{\epsilon_b}}$ | Relative latent period | 1 | 0.5 | 1 |
| $\delta_{\nu_b}$ | Relative probability of<br>detection | 1 | 1 | 0.1 |

**Table 2:** Summary of parameter values.

| Parameter | Meaning | Value | Reference |
| --- | --- | --- | --- |
| $1/\gamma$ | Length of the growing season | 120 days | Holt et al. (1999*a*); Rocco & Morabito (2016) |
| $\beta$ | Rate of secondary infection | $0.055/N \text{ day}^{-1} \text{ field}^{-1}$ | See main text |
| $\Delta$ | Time between roguing | 120 days | Illustrative |
| $\nu$ | Probability of detection | 1 | Illustrative |
| $\mu_U$ | Removal rate (unimproved) | $1/60 \text{ day}^{-1}$ | Illustrative |
| $\mu_R$ | Removal rate (resistant) | $1/60 \text{ day}^{-1}$ | Illustrative |
| $\mu_T$ | Removal rate (tolerant) | $1/1140 \text{ day}^{-1}$ | Illustrative |
| $\frac{1}{\epsilon}$ | Average latent period | 41 days | Holt et al. (1999*b*); Ber et al. (1990) |
| $\eta$ | Responsiveness of growers | 10 | Murray-Watson et al. (2022) |
| $Y$ | Maximum yield | 1 | All values scaled relative to yield |
| $L$ | Loss due to infection | 0.6 | Riley & Srinivasan (2019) |
| $\phi_T$ | Cost of tolerant crop | 0.1 | Fonsah et al. (2018) |
| $\phi_R$ | Cost of resistant crop | 0.1 | Fonsah et al. (2018) |
| $\phi_Q$ | Relative reduction in loss due to roguing | 0.7 | Illustrative |
| $N$ | Total number of fields/growers | 1 | Scaled to 1 |

Irrespective of the crop type planted in the field, and unless the crop in a field is rogued (see below), the season length (1/*γ*) is 120 days (Holt et al. (1999a), Rocco & Morabito (2016)). Replanting of any field occurs immediately after it is harvested. Susceptible fields become infected at rate *βI* day^−1^, where I is the proportion of infectious fields. Resistant crop, however, reduces the probability of infection per unit time by a factor *δ_β_R__* < 1. As tolerant crop does not reduce the probability of infection, we presume that *δ_β_T__* = 1. If fields are infected, they enter the “exposed” compartment where they remain latently infected (i.e. infected but not infectious) for an average of 1/*ϵ* days. During this time, we assume that they show no symptoms of infection and cannot infect other fields. We assume that resistant crops that become infected have a reduced rate of symptom development, increasing this latent period by a factor of *δ_ϵ_R__* < 1 (ensuring that *δ_ϵ_R__*/*ϵ* < 1/*ϵ*).

After becoming fully infectious and symptomatic, the ultimate fate of any field is to be either harvested or rogued. Roguing involves surveying fields (“scouting”) and removing any visibly-infected plants, and has been used as a means of TYLCV management for decades (Ioannu (1987), Polston et al. (1999), Polston & Lapidot (2007), Ddamulira et al. (2021)). As tomatoes can be harvested at different levels of maturity and ripen off-vine (Arah et al. (2015)), if there is infection in a field, growers may be better to harvest the entire field to prevent infection spreading. This will reduce the infection-associated yield loss by a factor *ϕ_Q_* such that the losses experienced by a grower who has rogued an infectious field are *ϕ_Q_L*). We assume scouting occurs at time intervals of Δ days and symptoms are detected with probability *v*. As tolerant crops reduce symptom severity, we presume that they have a lower probability of detection (reduced by multiplication by a factor of *δ_ν_T__* < 1) compared to unimproved and resistant crop (where *δ_νR_* = 1).

The removal rates of fields, *μ_b_* with *b* ∈ {*U, T, R*}, which represent such a program of roguing, are given by Cunniffe et al. (2014):

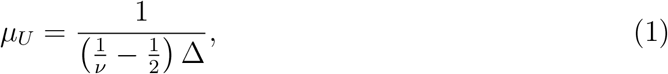

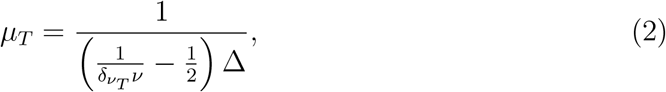

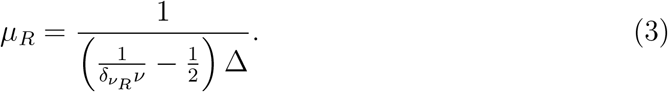

The epidemiological model is then given by:

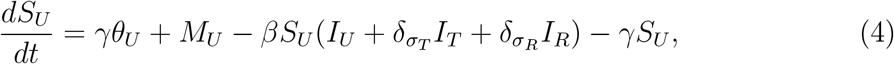

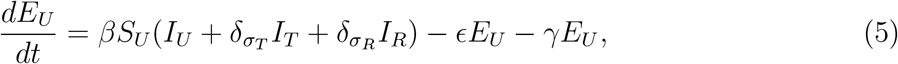

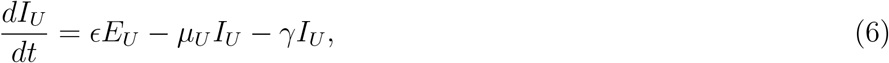

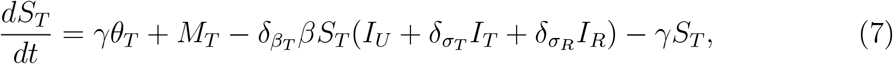

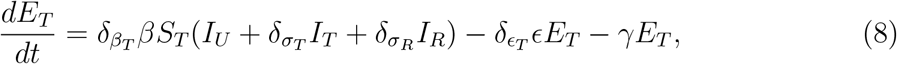

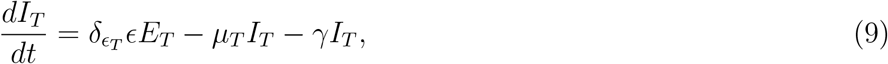

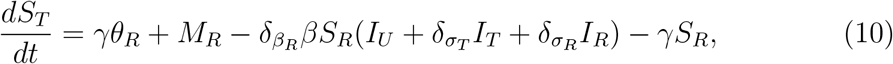

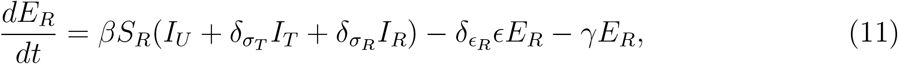

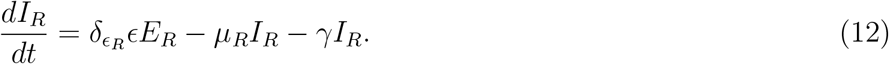

where terms *γθ_U_*, *γθ_T_* and *γθ_R_* are the rates of replanting for harvested fields, whilst *M_U_*, *M_R_* and *M_T_* are rates of replanting for rogued fields:

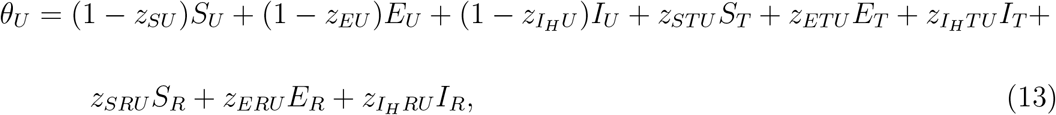

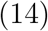

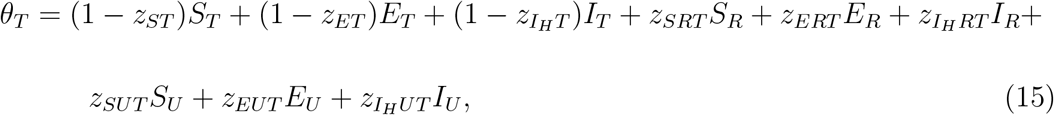

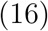

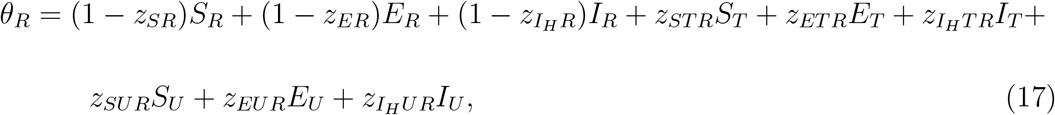

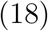

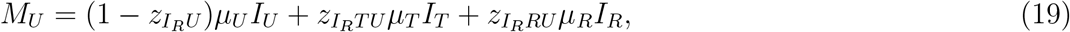

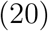

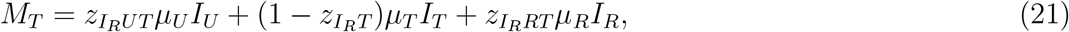

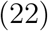

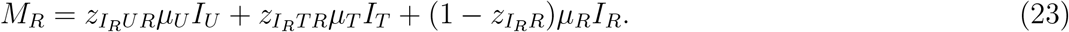

Note, a distinction is made between the replanting of rogued (*I_R_*) and harvested (*I_H_*) crop. For rogued unimproved fields, 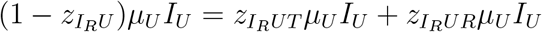 (i.e. the total rate at which rogued unimproved fields switch strategy must be equal to the rate at which they enter either other strategy, with equivalent expressions for tolerant and resistant crop). Similarly, for harvested unimproved fields, 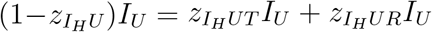. The exact forms of the *switching terms* (Murray-Watson et al. (2022)) is described below, including the meaning of the different number of subscripts (Equation 36).

36

Replanting in our model is represented by switching terms (Milne et al. (2016), McQuaid et al. (2017a), Murray-Watson et al. (2022)) which are the terms of the form *z_ab_* or *z_abc_* where *a* ∈ {*S,E,I_H_,I_R_*} and *b,c* ∈ {*U,T,R*} (note the additional complexity of our model means there is a change of notation compared to Murray-Watson et al. (2022)). The switching terms with three subscripts, *z_abc_*, represent the probability of a grower changing strategy from their current strategy *b* to strategy *c* based on their outcome, *a*, in the previous season (note that *b* ≠ *c*). For growers whose fields were susceptible (*S*) or exposed (*E*), the outcome depends solely on the epidemiological class of their field at the end of the season, whereas for growers whose fields were infectious, it depends also on whether that infectious field was rogued (subscript *R*) or harvested (subscript *H*). The two-subscript version, *z_ab_*, is the probability that a grower who harvested field type *a_b_* switches strategy. The forms of these terms are explained in Section 2.4 (below). A schematic of the model is shown in Figure 1.

**Figure 1:**
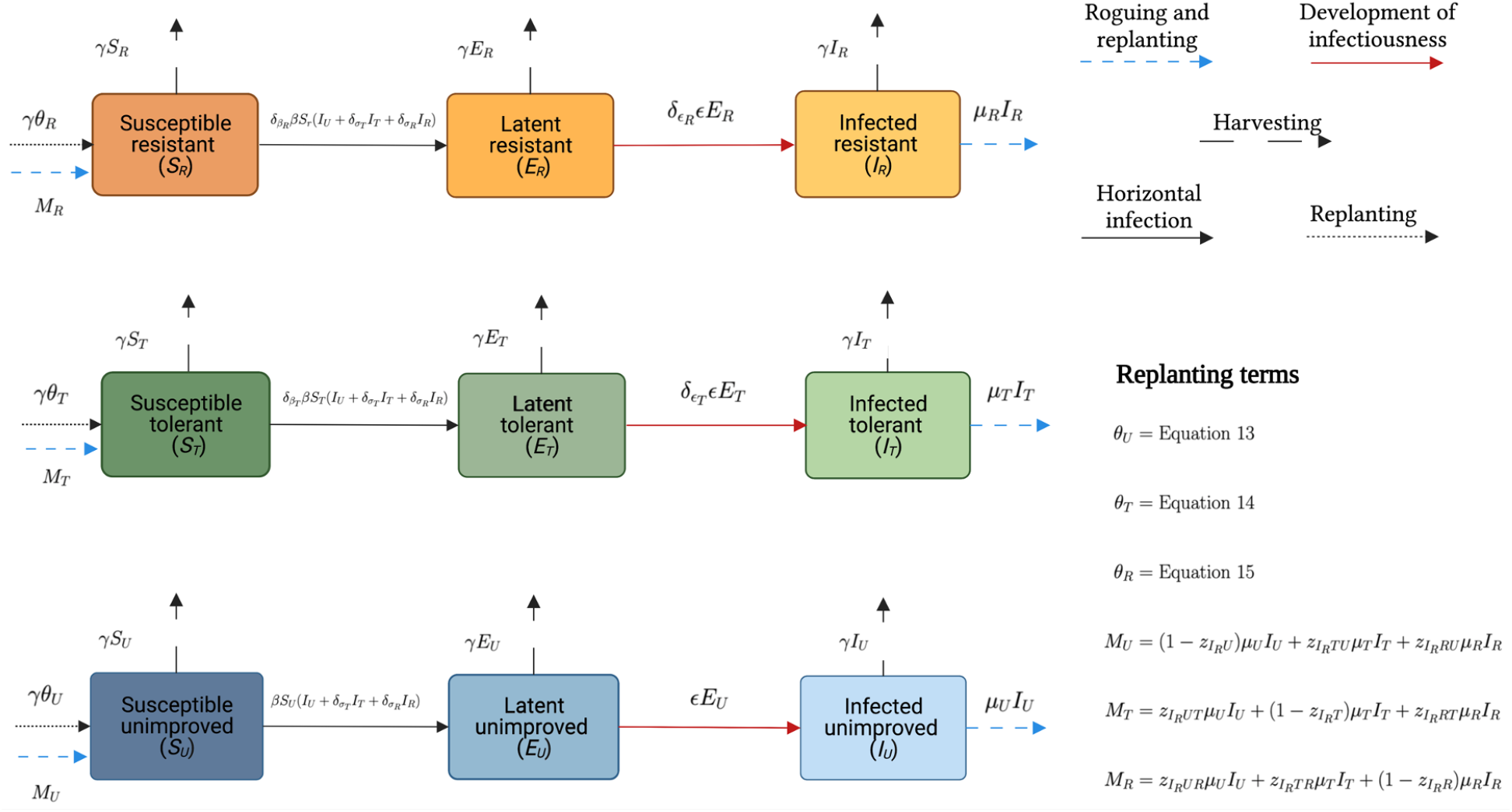
Schematic showing the structure of the model when growers can choose their strategy based on expected profits. We have three classes of growers; those who use unimproved seed (subscript *U*), those who use tolerant seed (subscript *T*), and those that use resistant seed (subscript *R*). The terms *θ_U_*, *θ_T_* and *θ_R_* are the rates of replanting for harvested fields, whilst *M_U_, M_R_* and *M_T_* are rates of replanting for rogued fields (Equations 13–23, with *M_U_* + *M_R_* + *M_T_* = *μ_U_I_U_* + *μ_R_I_R_* + *μ_T_I_T_*). Created with BioRender.com

### 3.2 Growers’ profits

The profit a grower earns is based on the control strategy that they previously used and their infection status at the time of harvest. Each crop type (*b* ∈ {*U,T,R*}) has specific associated costs and yield losses when infected with TYLCV.

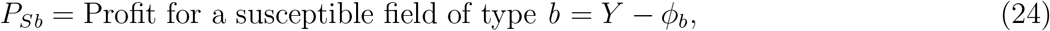

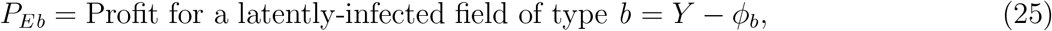

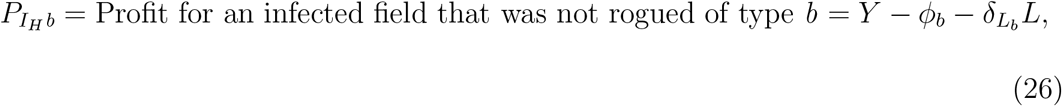

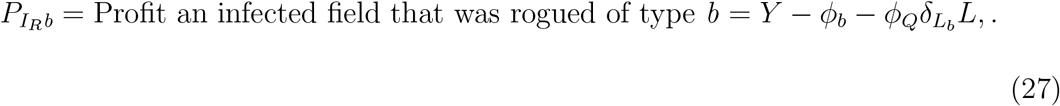

Notably, here, *ϕ_U_* = 0, since *ϕ_b_* represents the additional cost of using crop type b. For a full enumeration of the outcomes for growers, see Appendix 1.

For our parameterisation, we assume that latently-infected fields do not lose any yield, so *P_Sb_* = *P_Eb_, b* ∈ {*U,T, R*}. As growers who use unimproved crop and harvest susceptible or latently-infected fields pay no cost of control and sustain no yield loss, *P_SU_* = *P_EU_* is always the maximum achievable profit. The relative sizes of the remaining profits will depend on the values of the costs of the improved varieties (*ϕ_T_* and *ϕ_R_*), the infection-induced yield losses (*δ_L_L__L*, *δ_L_T__L* and *δ_L_R__L*) and the benefit of roguing (*ϕ_Q_*). Under the default parameterisation (Table 2), the ordering of the profits is given by:

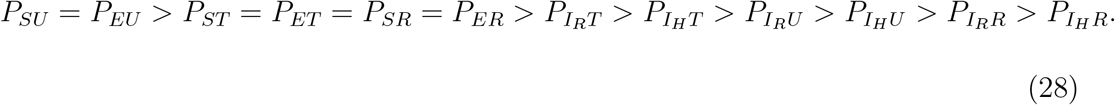

### 3.3 Calculating expected profits

The growers’ decision of which crop variety to use will depend on how each individual grower’s profit from the previous season compares to the expected profit of the alternative crop types. These expected profits are based on the probability of attaining each outcome outlined in Equations 42–53, which in turn depends on the probability of infection for each crop type. The complete derivation is provided in Appendix 1; the simplified expressions for the expected profits for unimproved (*P_U_*), tolerant (*P_T_*) and resistant (*P_R_*) are:

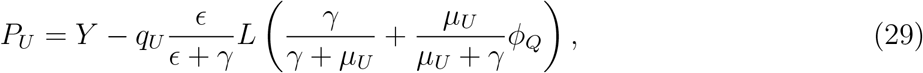

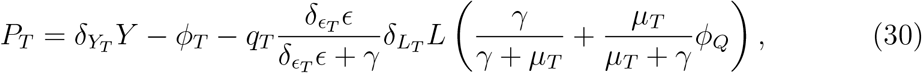

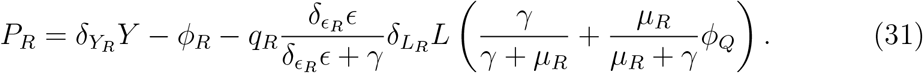

### 3.4 Switching terms

The switching terms (Murray-Watson et al. (2022)) affect the rate at which growers move from their current strategy to one of the alternative strategies.

In previous models (Milne et al. (2016), McQuaid et al. (2017a), Saikai et al. (2021) and Murray-Watson et al. (2022)), growers only had one alternative strategy with which to compare profits. Now, growers must consider two alternatives to their current strategy. Growers first assess the expected profits of the two alternative strategies and only compare their outcome with the highest expected profit of the two alternatives. If both alternatives have the same expected profit (so, for example, if the expected payoff for tolerance and resistance are the same (*P_T_* = *P_R_*)), and therefore the grower should have the same probability of switching into each strategy, growers chose to compare with the profit associated with the crop type most growers use. This follows from “descriptive norms”, where individuals follow what the majority of other people are doing (Cialdini et al. (1990), Sinclair & Agerström (2021), Lazić et al. (2021)). Additionally, there is some evidence that growers will be more likely to participate in control if other growers also participate (Milne et al. (2018)). If both the expected profits and the proportion of growers using each strategy are the same, then half of the growers changing strategy will go to each alternative.

In their general form, the switching terms have the following structure, where a grower with outcome *P_ab_*, contemplates switching into strategy *c* or *d, a* ∈ {*S, E, I_H_, I_R_*} and *b,c,d* ∈ {*U, T, R*},*b* = *c* = *d*.

If *P_c_* > *P_d_* or *P_c_* = *P_d_* and *C > D* (where *C* and *D* are the proportion using strategies *c* and d), then:

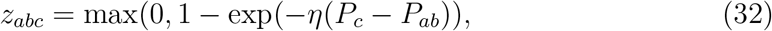

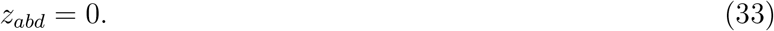

In the case where both the profits and the proportion of growers using the alternative strategy are equal, growers changing strategy will be divided evenly between the two alternative strategies. So, if *P_c_* = *P_d_* and *C* = *D*, then:

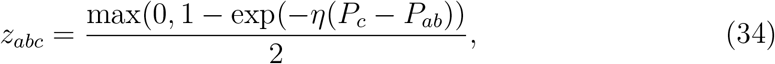

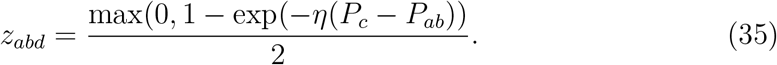

To simplify writing the model, we can then say:

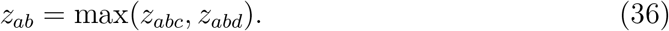

where *z_ab_* is the probability that a grower who harvested field of type *a_b_* leaves strategy *b* to adopt either strategy *c* or *d*.

### 3.5 Calculating the Pareto front and Gini coefficients

We use the concept of Pareto efficiency to dually optimise two conflicting objectives. Our first objective is to maximise the average profit of growers at equilibrium, calculated as the yield they achieve less any investment they have made to control for disease (based on the outcomes outlined in Equations 24–27). Our second objective is to minimise the spending of a central planning body which subsidises the cost of tolerant and resistant crops.

If the cost to the planner of the crop type is *ϕ_T,max_*, and the cost paid by growers is *ϕ_T_* then the subsidy to growers for the use of tolerant crop *σ_T_* = *ϕ_T,max_* – *ϕ_T_* (and, equivalently, for resistant crop is *σ_R_* = *ϕ_R,max_* – *ϕ_R_*). The total cost to the planner is then

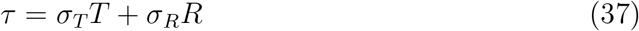

where *T* = *S_T_* + *E_T_* + *I_T_* and *R* = *S_R_* + *E_R_* + *I_R_* (i.e. are the proportions of growers using tolerant and resistant crop, respectively). We use Pareto optimality to establish the optimal allocation of subsidies between tolerant and resistant crop.

We first run the model for different values of *ϕ_T_* and *ϕ_R_* (400 values between 0 and 0.4 for each parameter) to find the equilibrium profit of growers and cost to the planner. We then use these two results as inputs to *get frontier ()* function from the KraljicMatrix package in R (Boehmke et al. (2017)), which calculates the Pareto front.

Though the Pareto front shows the optimal outcomes for a given set of parameters, one objective may be more heavily prioritised over the other. To quantify the degree of fairness between outcomes, we calculate the Gini coefficient (Equation 38). Originally developed as a measure of income inequality (Gini (1936)), the Gini coefficient calculates the extent to which a solution deviates from total equality between objectives. In that sense, the Gini coefficient is related to fairness but does not consider the optimality of the solution: a scenario with a low Gini coefficient (indicating a high degree of fairness between objectives, with a value of zero indicating perfect equality) may be a suboptimal solution for one or both objectives. Similarly, in some scenarios, the Gini coefficient may be small due to the number of objectives being considered: if just one individual’s objective is perfectly optimised, and all others are ignored, then *G* =1 – 1/*n*, where *n* is the number of objectives. The coefficient has been used to estimate regional inequalities in the risk of veterinary epidemics (Li et al. (2020)), geographical variability in STD incidence (Elliott et al. (2002)) and in ecology to evaluate management programmes and agricultural productivity (Li et al. (2017), Zaffaroni & Bevacqua (2022)). The Gini coefficient for a given scenario *i* is calculated by (Dorfman (1979), Zaffaroni & Bevacqua (2022)):

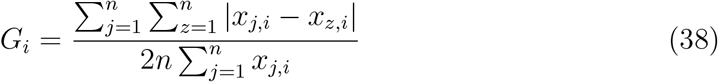

where *n* is the number of objectives considered and *x_j,i_* and *x_z,i_* are the values of the objectives for scenario *i*.

Calculating the Gini coefficient requires both objectives to be measured on the same scale, so we must aim to maximise or minimise both objectives. We therefore calculate the relative cost to the planner, *τ*\* as:

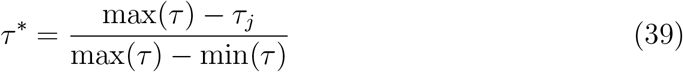

where max(*τ*) and min(*τ*) are the maximum and minimum costs to the planner that lie along the Pareto front and *τ_j_* is the cost value for a combination of *ϕ_R_* and *ϕ_T_*. This scales the cost to the planner between 0 and 1. Maximising this relative cost, *τ*\*, is equivalent to minimising the actual cost, *τ*.

Similarly, we must also rescale the profits so that they are normalised between 0 and 1:

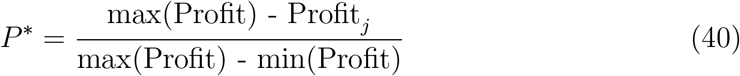

where max(Profit) and min(Profit) are the maximum and minimum profits to growers that lie along the Pareto front, Profit_*j*_ is the objective value for a particular combination of *ϕ_R_* and *ϕ_T_*.

We use the Gini coefficient to evaluate how a particular subsidy scheme benefits the planner relative to the growers, giving a measure of the fairness of each scheme. Since we have only two objectives, the Gini coefficient will be between 0 and 0.5.

## 4 Results

### 4.1 Equilibria for the three-strategy model

Using the next-generation method (NGM, van den Driessche (2017); Appendix 3) we find *R*_0_ for the model to be:

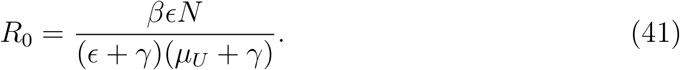

At the disease-free equilibrium, only unimproved crop is used by growers. The absence of disease makes control obsolete, so no growers should pay the cost of control. In Equation 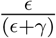 is the probability that an unimproved field will become infectious before it is harvested and 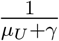 is the mean time in the *I_U_* compartment. The number of infections caused by these infectious fields is *βN*.

There are eight possible long-term outcomes for the model:

- **Disease-free equilibrium (DFE):** where *R*_0_ < 1 and no growers control for disease.
- **“No control” equilibrium:** where disease is endemic but no growers use improved crop (*N* = *U*).
- **“All tolerant” equilibrium:** where disease is endemic and growers only use tolerant crop (*N* = *T*).
- **“All resistant” equilibrium:** where disease is endemic and growers only use resistant crop (*N* = *R*).
- **Three-strategy equilibrium:** where disease is endemic and crops of all three varieties are in use (*N* = *U* + *T* + *R*).
- **“Tolerant and unimproved” equilibrium:** where disease is endemic and growers use either tolerant or unimproved crop (*N* = *U* + *T*).
- **“Resistant and unimproved” equilibrium:** where disease is endemic and growers use either tolerant or unimproved crop (*N* = *U* + *R*).
- **“Tolerant and resistant” equilibrium:** where disease is endemic and growers use either tolerant or resistant crop (*N* = *T* + *R*).

Additionally, the model presented in Murray-Watson & Cunniffe (2022) allowed for a bistable region when *R*_0_ < 1. To prevent this from affecting our results, we always begin the models with the initial conditions outlined in Table 3, which guarantees a disease-free equilibrium in the bistable region for the default parameterisation.

**Table 3:**
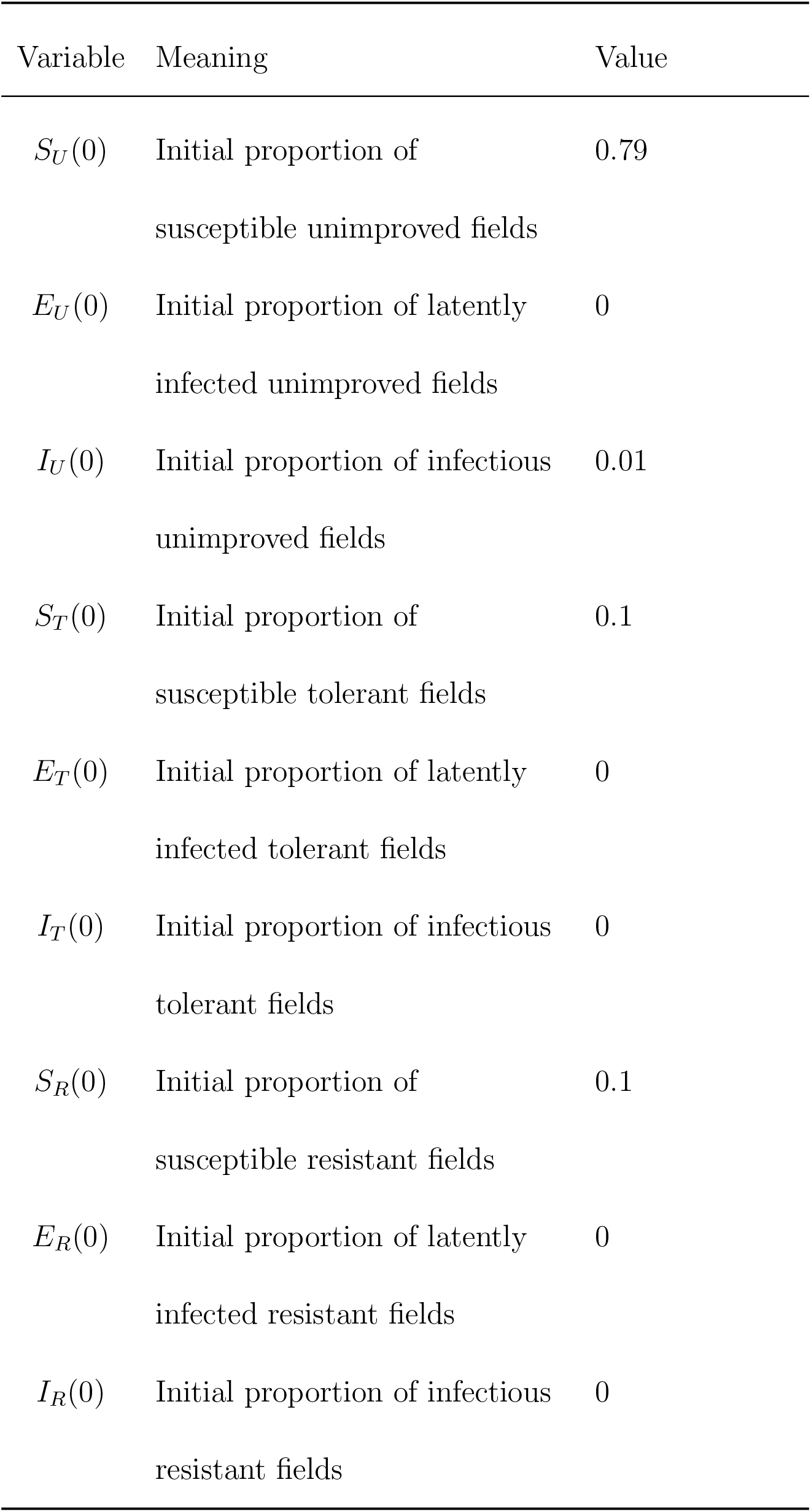
Default initial conditions.

| Variable | Meaning | Value |
| --- | --- | --- |
| $S_U(0)$ | Initial proportion of<br>susceptible unimproved fields | 0.79 |
| $E_U(0)$ | Initial proportion of latently<br>infected unimproved fields | 0 |
| $I_U(0)$ | Initial proportion of infectious<br>unimproved fields | 0.01 |
| $S_T(0)$ | Initial proportion of<br>susceptible tolerant fields | 0.1 |
| $E_T(0)$ | Initial proportion of latently<br>infected tolerant fields | 0 |
| $I_T(0)$ | Initial proportion of infectious<br>tolerant fields | 0 |
| $S_R(0)$ | Initial proportion of<br>susceptible resistant fields | 0.1 |
| $E_R(0)$ | Initial proportion of latently<br>infected resistant fields | 0 |
| $I_R(0)$ | Initial proportion of infectious<br>resistant fields | 0 |

Parameterisations that lead to the disease-free, all-tolerant, tolerant and resistant, unimproved and resistant, unimproved and tolerant, and three-strategy equilibria are shown in Figure 2, in which we vary the rate of horizontal transmission (*β*) and in (A) the cost of both improved crops (*ϕ_R_* and *ϕ_T_*) and in (B) just *ϕ_T_* (*ϕ_R_* is fixed at 0.06 for demonstrative purposes). However, the externalities generated by both tolerant and resistant crop mean that some of these potential equilibria are only realised within a narrow range of parameter values (namely the “no control” equilibrium). The “all resistant” equilibrium is not possible, as those using unimproved crop can free ride off the efforts of those who plant resistant crop (Murray-Watson & Cunniffe (2022)).

**Figure 2:**
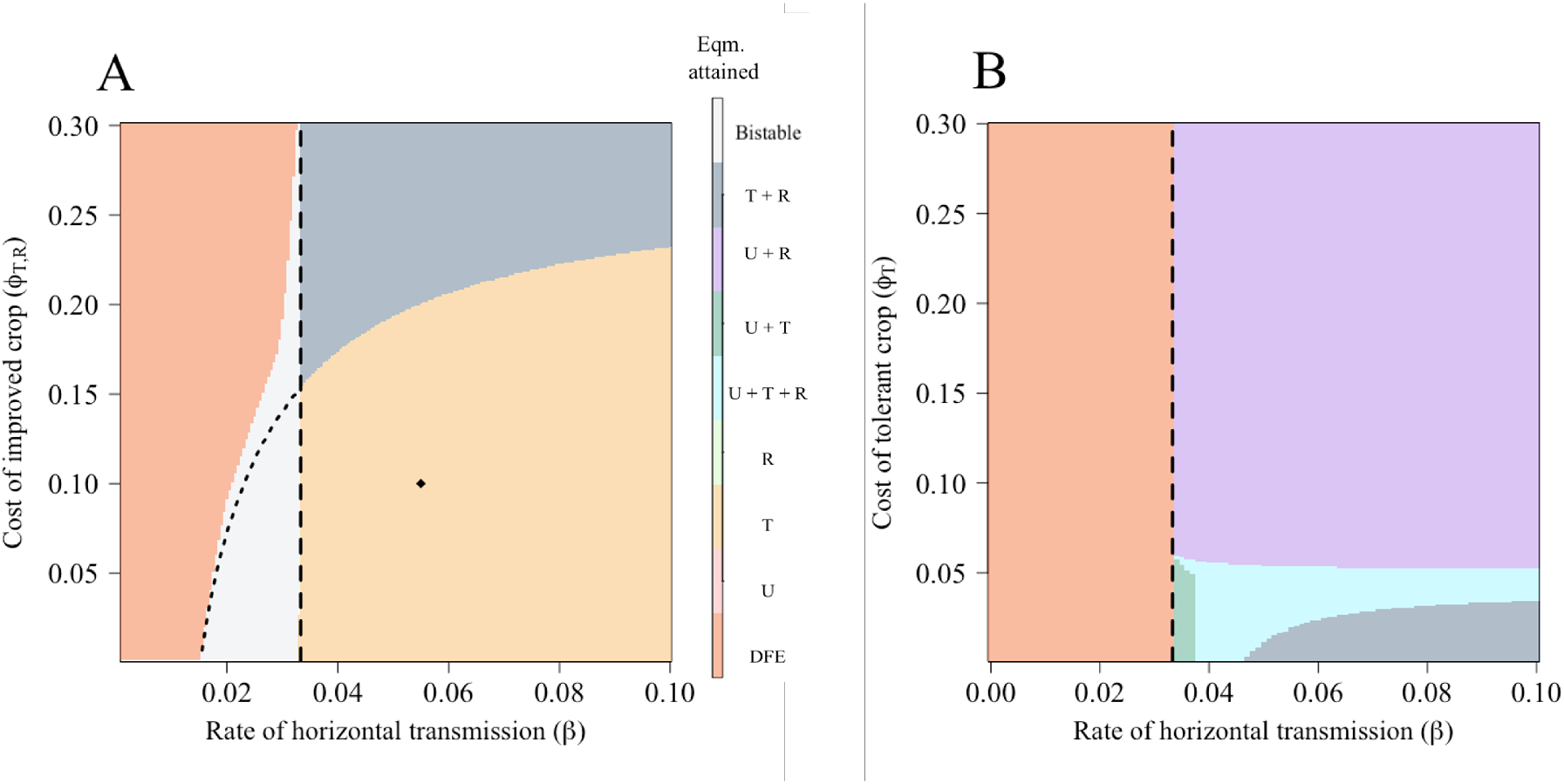
Effect of the cost of tolerant or resistant crop and the rate of horizontal transmission on the equilibrium values. (A) Uses the default parameterisation, whilst (B) has *ϕ_R_* = 0.06, *δ_ν_T__* = 1 (as might be the case if disease detection was done via a genetic test), *δ_L_T__* = 0.3 and *δ_β_R__* = 0.1. In both cases, a disease-free equilibrium persists once *R*_0_ < 1, though there is a bistable region shown in light grey between the limits of *β_U_* = 0.020 and 0.0333 day^−1^ in (A). The equilibrium realised within this region is either a mixed tolerant and unimproved crop equilibrium or an all-tolerant equilibrium (indicated by the dotted line). In (B), lowering the costs and losses associated with resistant crop allowed for a mixed equilibrium between resistant and unimproved crop and a three-strategy mixed equilibrium between all three crop types. The dashed vertical lines show *R*_0_ = 1.

Which long-term outcomes are possible depends strongly on parameter values, particularly the parameterisation of resistance and tolerance. When the cost of both improved crop varieties varied (Figure 2(A)), lower costs incentivised the use of tolerant crop, and as the rate of horizontal transmission (*β*) increased, more growers used tolerant crop. The high probability of infection meant that growers would be better off using tolerant crop and limiting the damage due to disease (as, under this parameterisation, the losses in tolerant crop were a tenth of those for resistant or unimproved crop). As the benefits of tolerant crop are felt privately by those who use them, there is more incentive to use these varieties and an “all-tolerant” equilibrium is possible even at relatively high costs of control.

If we keep the cost of resistant crop fixed at a low price (*ϕ_R_* = 0.06), and parameterise it such that it is significantly less susceptible to infection (*δ_β_R__* = 0.1), its use is much more widespread (Figure 2(B)). At low values of *ϕ_T_*, mixed equilibria with tolerant crop are achieved, particularly at higher values of *β* when infection becomes more likely to occur. However, under this parameterisation, the tolerant crop is both more easily detected via roguing (*δ_ν_R__* = 1) and is less effective at limiting yield loss (*δ_L_* = 0.3), so there is less of an incentive for growers to use it compared to the parameterisation in Figure 2(A). As *ϕ_T_* increases, growers stop using tolerant crop and instead a larger proportion “free-ride” off of the efforts of those that use resistant crop. As resistant crop generates positive externalities for other growers, the fields of the non-controllers have a reduced infection pressure without incurring any costs. Consequently, an “all-resistant” equilibrium is not reached.

The bistable region between *β* = 0.02 and 0.0333 day^−1^ in Figure 2(A) varies between either the DFE and “all tolerant” equilibria or the DFE and “tolerant and resistant” equilibria depending on parameter values, with the dotted line showing the distinction between these equilibria. As in Murray-Watson & Cunniffe (2022), a high initial proportion of tolerant crops (or infectious crops) causes the system to go to the disease-endemic equilibria.

### 4.2 Effect of parameters relating to tolerance and resistance on behaviour

The primary characteristic of tolerant crop is its ability to limit yield loss if infection occurs, so the losses experienced by growers of tolerant crop are lower than those of unimproved or resistant crop (*δ_L_T__L* < *L*). At low values of *δ_L_T__* (Figure 3), many growers therefore use tolerant crop. As tolerant crop is less likely to be rogued, infection builds up, further incentivising growers to use tolerant crop.

**Figure 3:**
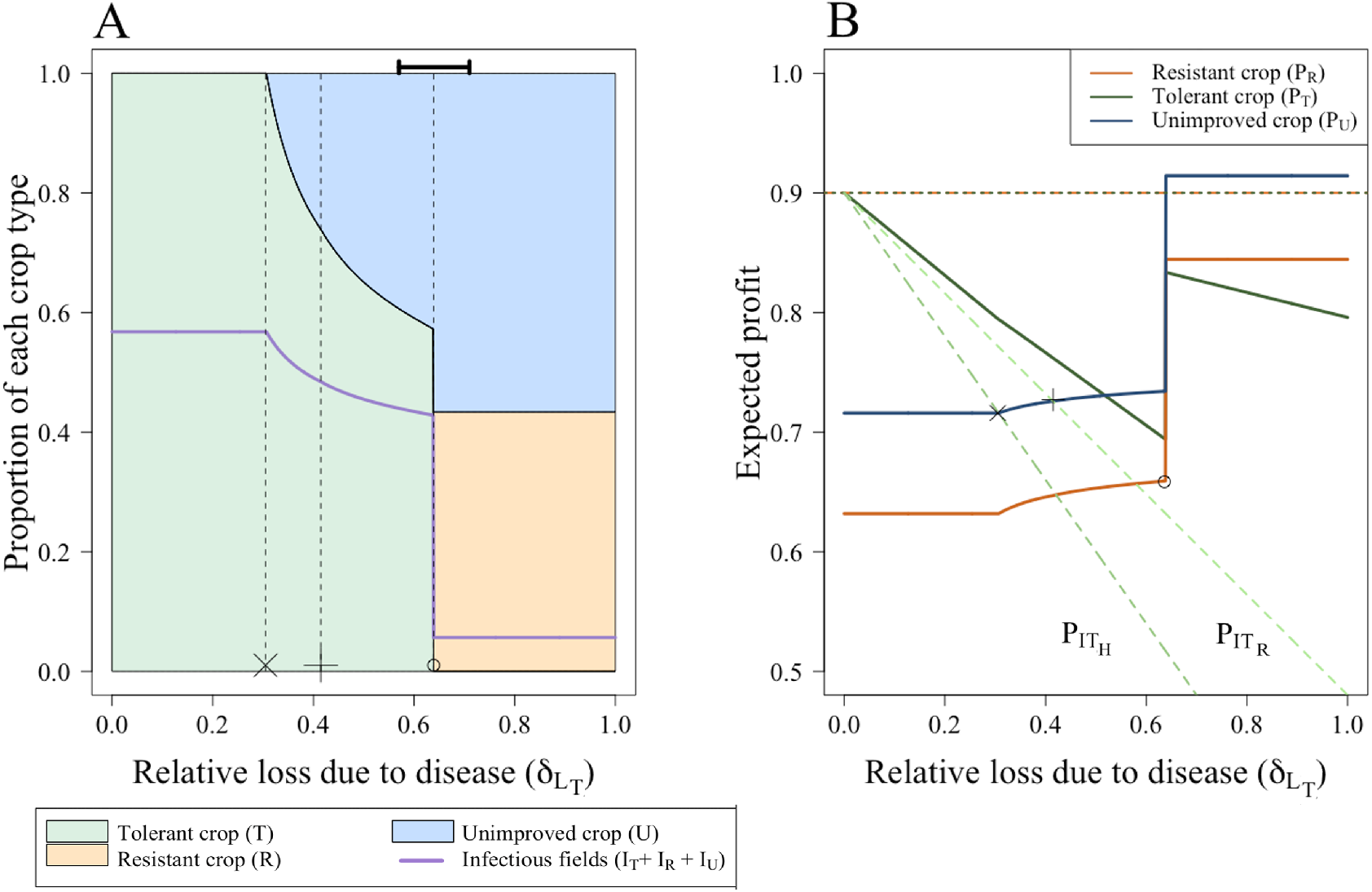
Effect of the relative loss due to infection on the choice of unimproved, tolerant or resistant crop. (A) The proportion of infectious fields (*I_T_* + *I_R_*), tolerant (*T*) and resistant (*R*) fields. (B) The expected profit for growers using unimproved, resistant or tolerant crop. The dashed orange and red lines show the profit for susceptible and latently-infected tolerant and resistant fields (*P_ST_* = *P_ET_* = *P_SR_* = *P_ER_* = 0.9), and the green dashed lines show the profits for a rogued and harvested infected tolerant field. For low values of *δ_L_T__* (such that *δ_L_T_L_* << *δ_L_R__L,δ_L_T__L* << *L*), all growers use the tolerant crop. However, once *δL__T__* > 0.31, growers who have harvested infected tolerant crop should have a non-zero probability of switching to unimproved crop. Similarly, once *δ_L_T__* > 0.41, growers who have rogued tolerant crop should also have a non-zero probability of switching to unimproved crop. Once *δ_L_T__* > 0.64 (when *PT* = *PR*), all non-controllers who harvested infected crop should consider switching to resistant crop, rather than tolerant crop. This causes a decrease in those using tolerant crop and an increase in those planting resistant, which also causes a decrease in the proportion of infected fields (A). The black bar in (A) shows the region where bistability affects the switch between equilibria. Save for *δ_L_T__*, parameters and initial conditions are as in Table 2 and Table 3 respectively.

As *δ_L_T__* increases, the value of the expected profit of those growing tolerant crop (*P_T_*) falls. For this parameterisation, at *δ_L_T__* ≈ 0.31 (“X”), the profits of growers who harvested infected tolerant crop are lower than the expected profits of non-controllers (*P_I_H_T_* < *P_U_*), so controllers that have harvested infected crop have a non-zero probability of switching strategy. As the expected profit for unimproved crop is higher than that of resistant crop (*P_U_* > *P_R_*), these growers only consider switching to unimproved crop. Similarly, at *δ_L_T__* ≈ 0.405 (“+”), *P_I_R_T_* < *P_U_* and growers of tolerant crop who have rogued their fields should also consider switching to unimproved crop. These changes to the switching terms cause sharp changes in the proportion of growers using each strategy.

At the value of *δ_L_T__* ≈ 0.64 (“O”), growers of tolerant crop who have rogued their infected fields earn less than the expected profit of those using resistant crop (*P_I_R_T_* < *P_R_*). These growers start switching strategy, causing a decrease in the proportion using tolerant crop and an increase in the expected profits of both tolerant and resistant crop. At *δ_L_T__* ≈ 0.64 *P_T_* = *P_R_*: the model set-up means that growers of unimproved crop who harvested infected fields compare with the profit of whichever strategy has more growers using it, which, in this case, is tolerant crop. Once *δ_L_T__* > 0.64 (“O”), *P_T_* < *P_R_* and the growers of infected unimproved crop now consider switching to resistant crop. This causes a sharp increase in those using resistant crop and decrease in proportion of infected fields (Figure 3 (A) at *δ_L_T__* > 0.64). All expected profits rise with this decrease in infection (Figure 3 (B)), though the relative ordering of the profits does not change (*P_U_* > *P_R_* > *P_T_*). The response flattens out after *δ_L_T__* > 0.64 as no growers are using the tolerant crop, so no growers are affected by the changes in this parameter. The model will therefore run to the same equilibrium irrespective of further changes to *δ_L_T__*.

Between the values of *δ_L_T__* ≈ 0.51 and 0.71 (denoted by the black bar in Figure 3 (A)), there is bistability which changes the precise value of *δ_L_T__* at which growers stop using tolerant crop (“O”), as a function of the initial conditions of the model (Appendix 3).

When considering the efficacy of resistant plants, only when the relative susceptibility of resistant plants (*δ_β_R__*) is low is there resistant crop at equilibrium. Once the resistant crop is more susceptible to infection, growers prefer to use the tolerant crop and sustain lower losses once infection occurs. For the parameterisation in Table 2, and as *δ_β_R__* varies, when *δ_β_R__* = 0.08, all growers use tolerant crop (“all tolerant” equilibrium), so the responses to changes in *δ_β_R__* flatten out as there are no resistant fields with reduced susceptibility. The remaining parameters are unchanged, so the model goes to the same equilibrium as parameter *δ_β_R__* is varied.

Similarly to Figure 3, between the values of *δ_β_R__* = 0.054 and 0.077 (denoted by the black bar in Figure 4), there is bistability which changes the point at which growers stop using resistant crop and only use tolerant crop (Appendix 3).

**Figure 4:**
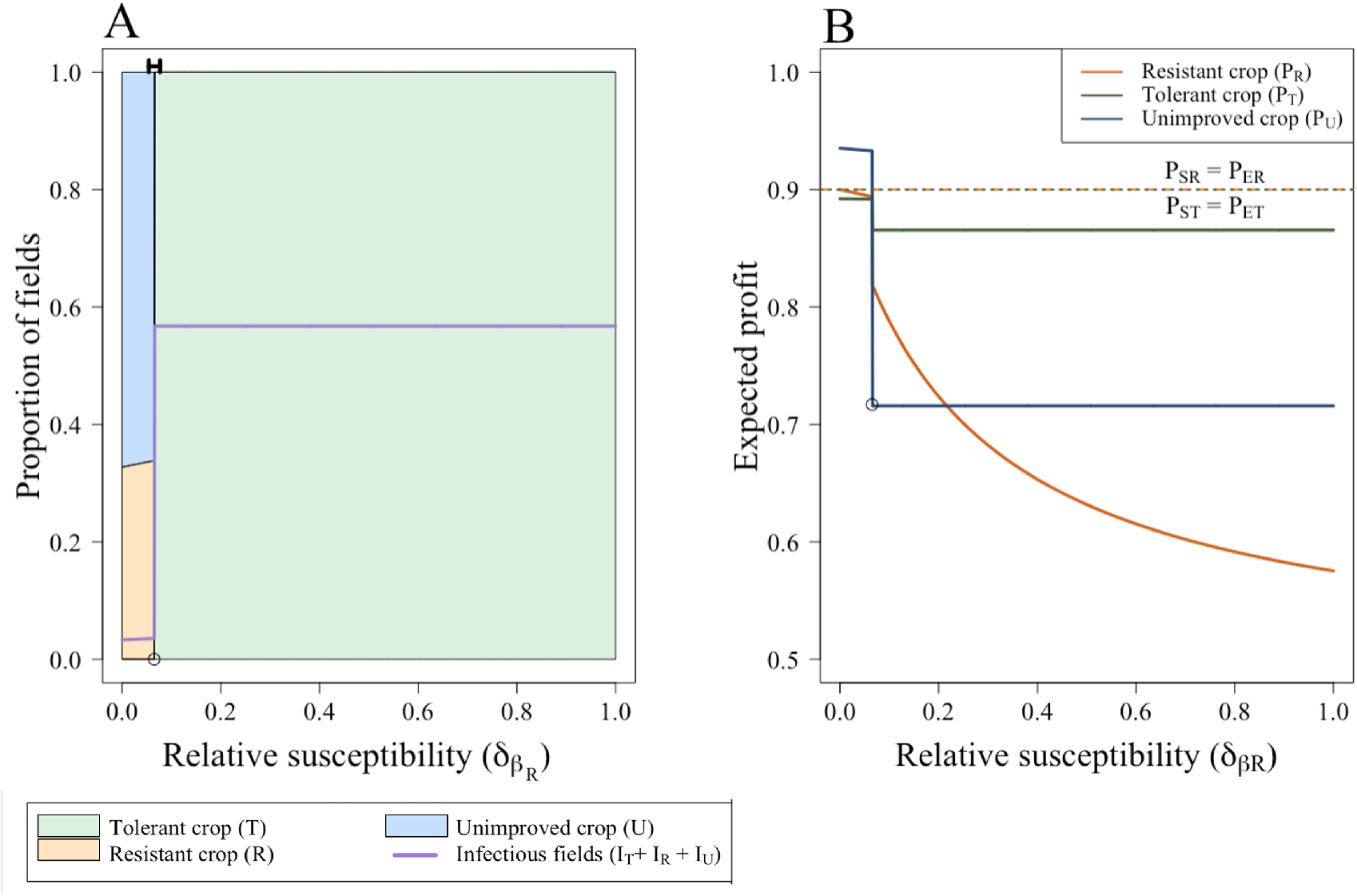
Effect of the relative susceptibility to infection on the choice of unimproved, tolerant or resistant crop. (A) The proportion of infectious fields (*I_T_* + *I_R_* + *I_U_*), tolerant (*T*) and resistant (*R*) fields. (B) The expected profit for growers using unimproved, resistant or tolerant crop. The dashed orange and red lines show the profit for susceptible and latently-infected tolerant and resistant fields (*P_ST_* = *P_ET_* = *P_SR_* = *P_ER_* = 0.9). Most growers use the resistant crop for low values of *δ_β_R__*. The remaining growers free-ride off of these actions and do not use any improved crop. However, as *δ_β_R__* increases, the benefits to growers of resistant crop are lower, and growers start using tolerant crop. This causes an increase in the proportion of infected fields. Save for *δ_β_R__*, parameters and initial conditions are as in Table 2 and Table 3 respectively.

### 4.3 Pareto optimality

Here, as an example, we calculate the Pareto front when neither improved crop type is very effective (the relative susceptibility of resistant crop is *δ_β_R__* = 0.5 and the relative loss due to disease in tolerant crop is *δ_L_T__* = 0.5). Figure 6 shows the Pareto front when the planner is trying to minimise the cost of the subsidy scheme whilst also maximising the average profit of growers, with both quantities considered at the eventual equilibrium of the model.

**Figure 6:**
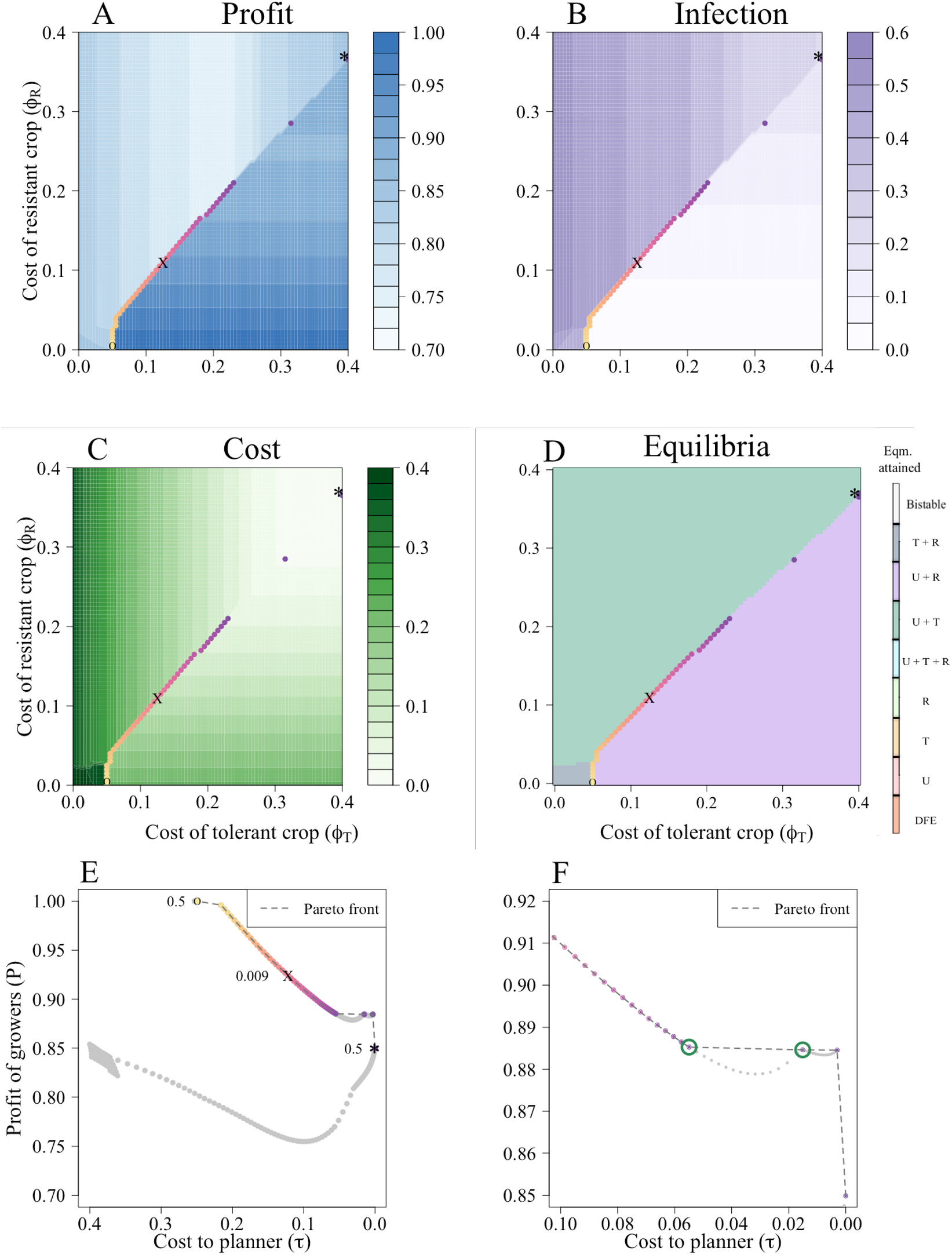
Pareto front when both the tolerant and resistant crop are only moderately effective. (A) The profit to growers. (B) The level of infection. (C) The cost to the planner (*τ*, Equation 37) and (D) which equilibria are attained. The dots in (A) - (D) represent the combination of *ϕ_T_* and *ϕ_R_* that lie along the Pareto front in (E), and all are parameter combinations that disincentivise the use of tolerant crop. (E) Pareto front when both the tolerant and resistant crop are only moderately effective. The individual grey dots correspond to all pairs of values of (*ϕ_T_, ϕ_R_*) considered in our two way scan; the coloured dots are those lying on the Pareto front. The darker the dots, the more the low costs to the planner have been prioritised. The Gini coefficients are shown for the fairest scenario (*G* = 0.009, when *τ* = 0.13 and *P* = 0.93; “X” on the graphs) and the least fair scenarios (*G* = 0.5; “O” when the profit is prioritised and “*” when the costs to the planner are). (F) Pareto dominance between *τ* = 0.06 and *τ* = 0.0, which corresponds to the “break” in the Pareto front in (E).

We plotted the Pareto front onto a two-way scan of the cost of resistant and tolerant crop (*ϕ_R_* and *ϕ_T_* respectively) to investigate the subsidy regime that produces optimal outcomes (Figure 6(A)-(D)). The strategies that guarantee the best outcome for both the growers and the planner are to use any combination of *ϕ_R_* and *ϕ_T_* that discourages widespread use of tolerant crop (Figure 6(D), where the Pareto front lies along the edge of the “U + T” and the “U + R” equilibrium). Below the value of *ϕ_R_* = 0.365, the optimal subsidy schemes ensure a mixed equilibrium of unimproved and resistant crop (Figure 6(D)). Above this value (for which the only optimal strategies are *ϕ_T_* = 0.4, *ϕ_R_* = 0.365 or *ϕ_T_* = 0.4, *ϕ_R_* = 0.367) resistant crop is too expensive and growers instead use tolerant crop, though it is always at relatively low levels (around 10% of fields are planted with tolerant crop, Appendix 3 Figure 4).

Though the use of tolerant crop earns the growers high profits, the resulting increase in infection pressure induces a positive feedback loop that incentivises other growers to also use tolerant crop. The planner will therefore have to subsidise more growers, so it is ultimately not economical for the planner, and no Pareto-optimal strategy lies in the region where there is high use of tolerant crop. Conversely, as resistant crop provides benefit to growers who do not use it (i.e. “free ride” off the efforts of others), by subsidising a minority of growers to use resistant crop the planner can achieve good profit outcomes averaged over the population as a whole for lower costs.

Though the Pareto front lies along the diagonal, any points in a given row along to the right of the diagonal have the same value (as there is no tolerant crop at equilibrium, so changing the cost of tolerant crop has no effect on the equilibrium achieved; equivalently, all points in a given column above the diagonal have the same value as there is no resistant crop at equilibrium). They are, therefore, all considered to be Pareto optimal.

We then plotted the Pareto front for each combination of costs and profits from Figure 6(A) and (C) (Figure 6(E)). The grey dots are subsidy strategies that are dominated by other strategies that lie on the Pareto front. The Pareto front is broken between *τ* = 0.058 and *τ* = 0.015 (indicated by green circles in Figure 6(F)). These combinations of costs and profits are dominated by *τ* = 0.015 and *P* = 0.88, where the profit is at a local maximum. The profits earned by growers when 0.058 > *τ* > 0.016 are all *P* < 0.88, so both growers and planners can achieve better outcomes at *τ* = 0.015. Between 0.058 > *τ* > 0.016, the subsidisation scheme reduces use of the resistant crop, decreasing profits for the growers as more fields become infected.

The degree of subsidisation (and associated outcome) will depend on the planner’s weighting of the relative importance of the cost to the planner and the profit of growers. This can be measured using Gini coefficients. The Gini coefficients are shown for the extremes of equality: the least fair scenarios, with a Gini coefficient of 0.5, lie at either end of the Pareto front. Time courses for these different scenarios are shown in Figure 3 in Appendix 4.

When the profit to growers is 1.00 and the cost to the planner is 0.25, corresponding to *ϕ_R_* = 0.0 and *ϕ_T_* = 0.05 (marked “O” in Figure 6), a relatively high proportion of growers use resistant crop (≈ 49.8%, Figure 4(B) in Appendix 4). At the other end of the front, the profit to the growers is 0.85 and τ = 0. As *ϕ_R_* = 0.4 and *ϕ_T_* = 0.375 ((marked “*” in Figure 6)), so improved crop is effectively not sub-sidised. This results in no users of resistant crop and only around 10% of growers use tolerant crop (Figure 3(C) in Appendix 4), reducing the costs to the planner (this is also the only point on the Pareto front where any growers use tolerant crop; Figure 4(A) in Appendix 4).

The fairest scenario (*G* = 0.009, “X”) occurs when *ϕ_T_* = 0.12 and *ϕ_R_* = 0.105 (*τ* = 0.13 and *P* = 0.93). At this point, ≈ 40.3% of growers use resistant crop, none use tolerant crop (Figure 4 in Appendix 4), and only 6% of fields are infected (Figure 6 (B)). In this “U + R” equilibrium, the majority of growers can “free ride” off the efforts of those using resistant crop.

### 4.4 Effect of time on the Pareto front

Both the costs to the planner and the profits of growers depend on the proportion of fields of each type in the system. This has a strong temporal component; consequently, the Pareto front may change depending on the time horizon examined. We investigated this temporal effect after 1, 2, 3, 4, and 5 seasons, fixing the cost of resistant crop (*ϕ_R_*) to 0.1 for ease of analysis (that is, in this section, we only consider changes to the cost of tolerant crop, *ϕ_T_*).

At short time points (one or two seasons), nearly every subsidisation scheme lies on the Pareto front (Figure 7(A)-(D)). Each of these value of *ϕ_T_* are considered equally efficient, and produce a Pareto-optimal combination of outcomes for both the growers and the planners.

**Figure 7:**
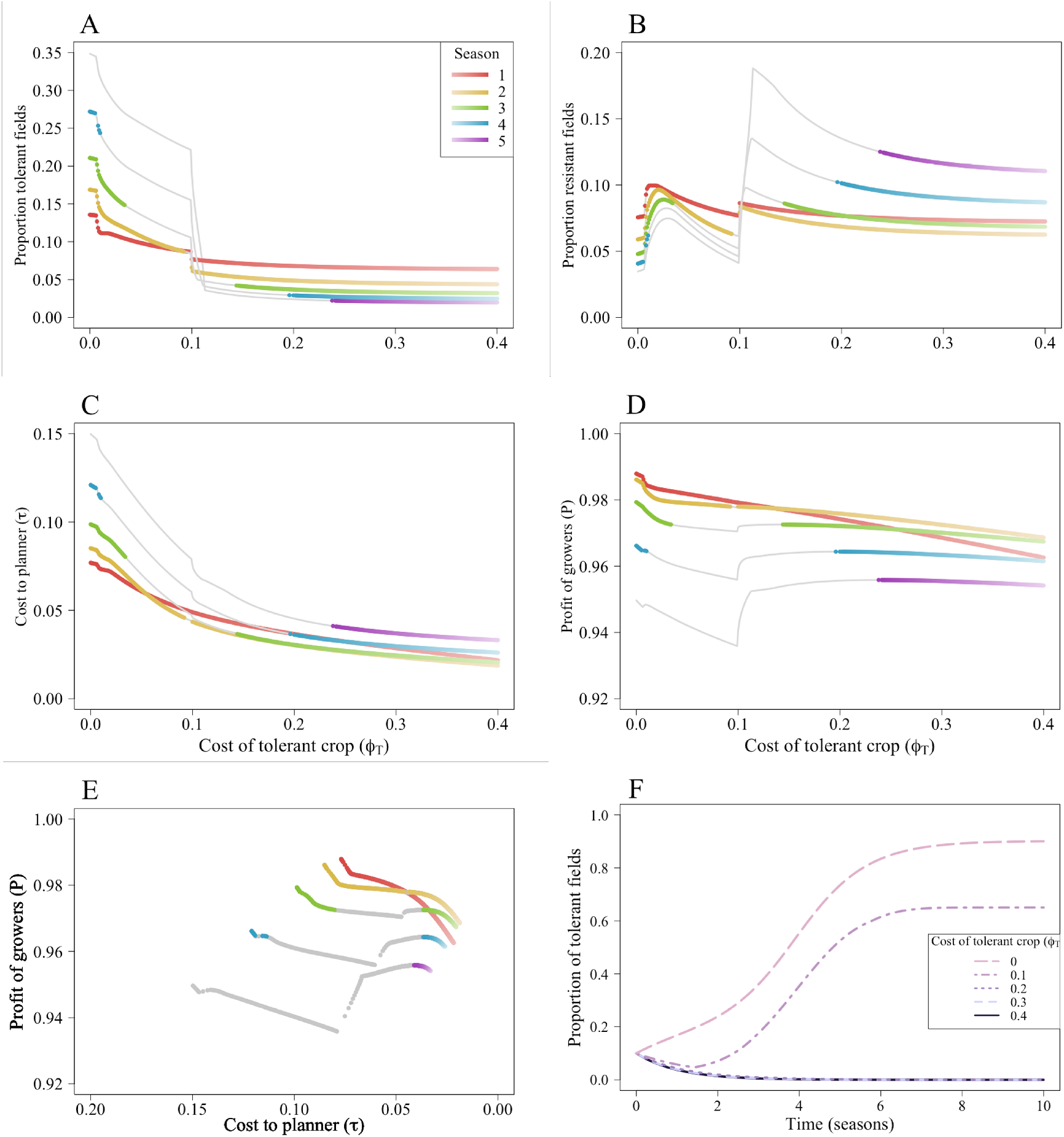
Pareto front at early seasonal time points when both the tolerant and resistant crop are only moderately effective. (A) The proportion of tolerant fields. (B) The proportion of resistant fields. (C) The cost of the subsidisation scheme to the planner. (D) The profit to the growers. The darker the colour, the more the profit of growers is prioritised over the cost to planners. Generally, the Pareto front lies along values of *ϕ_T_* that reduce the proportion of growers using tolerant crop and increase those using resistant crop. (E) At low time points, all all subsidisation schemes are Pareto-optimal, though as time progresses there are a narrower range of optimal solutions. (F) Once *ϕ_T_* > 0.2, no growers use tolerant crop after 5 seasons. Other than *δ_L_T__* = 0.5 and *δ_β_R__* = 0.5, parameters are as in Table 2. Note, in each of these graphs, the cost of resistant crop (*ϕ_R_*) is 0.1.

As the epidemic progresses (seasons three, four, and five), only higher values of *ϕ_T_* give Pareto-optimal solutions (Figure 7(A)-(D)). The higher cost of tolerant crop disincentivises its use, so fewer growers use it (Figure 7(A)). Though there is an increase in the proportion using resistant crop, uptake of resistant crop is still relatively low (Figure 7(B)) due to the ability of non-controllers to free-ride. Overall, the higher values of *ϕ_T_* lowers the costs to the planner, whilst maintaining high profits for the growers (Figure 7(C)-(E)).

Yet, between three and four seasons into the epidemic, low values of *ϕ_T_* also permit Pareto-optimal solutions. At these values, relatively few of the growers are using tolerant crop (Figure 7(F)) and the costs to the planner are relatively low, whilst the profits to the grower are high (Figure 7(C)-(D)).

The Pareto fronts show that, over these shorter time horizons, there is often a small difference in the profits and costs for the optimal cases (Figure 7(E)). We again find that the costs of improved crop that discourage the use of tolerant crop are optimal. For each time horizon, the Pareto optimum is achieved at higher values of the cost of tolerant crop to growers (*ϕ_T_*), which discourages the use of tolerant crop. This implies that in the early stage of an epidemic, there is not as severe a trade-off between the optimal outcomes for the planner and the grower.

Figure 7(F) shows a time course of the proportion of growers using tolerant crop. At low values of the cost of tolerant crop (*ϕ_T_*), which are only part of the Pareto-optimal sets for short time horizons (between one and four seasons), tolerance does not die out. At higher values, tolerance dies out within five seasons. Therefore, whilst lower values of ϕT can sometimes provide Pareto-optimal solutions, higher values give optimal outcomes irrespective of the time horizon.

The same results hold for other parameterisations of tolerance and resistance - irrespective of how good or bad the improved crop varieties are, the Pareto optimal strategies are those in which the use of tolerant crop is discouraged.

## 5 Discussion

Human behaviour has increasingly been investigated in the context of plant disease epidemiology (Milne et al. (2016), McQuaid et al. (2017a), Saikai et al. (2021), Murray-Watson et al. (2022), Murray-Watson & Cunniffe (2022), Milne et al. (2020), Bate et al. (2021)), where models allow growers choose between engaging in disease control or taking no action. The proportion of growers using control depends on the control mechanism itself, with previous studies investigating the uptake of clean seed systems amongst cassava growers (McQuaid et al. (2017a), Murray-Watson et al. (2022)), transgenic herbicide-resistant maize (Milne et al. (2016), Saikai et al. (2021)) or disease-tolerant or -resistant crop (Murray-Watson & Cunniffe (2022)). Uptake will also depend on the relative costs of control and losses due to disease, amongst other factors not investigated here (such as the social connections between growers). However, in each of these examples, growers could only choose whether to use control or not, not between different types of control.

When investigating the effect of having crop that was either tolerant or resistant to Tomato Yellow Leaf Curl Virus (TYLCV) available to growers (alongside unimproved crop), Murray-Watson & Cunniffe (2022) found that widespread use of tolerant crop was often achieved. Tolerant crop only benefited those growers using it, and due to the lower rate of removal via roguing, they increased the infection pressure on other growers (generating negative externalities). However, when resistant crop was available, such high levels of adoption were not achieved. Resistant crop protected other, unimproved fields (generating positive externalities), who can *free-ride* off the efforts of those using resistant crop.

These results were corroborated in the work presented here. Even when growers have a choice between three crop types (unimproved, disease-tolerant or -resistant), the positive feedback loop induced by the use of tolerant crop is sufficient to ensure that growers will overwhelmingly use tolerant crop, even when tolerant crop was relatively ineffective (Figure 3(A)). This positive feedback loop means that tolerant crop is a “strategic complement” (a strategy that incentivises others to adopt the strategy as more individuals use that strategy; Hennessy (2008), Murray (2014), Delabouglise & Boni (2020)). Resistant crop, by contrast, is a “strategic substitute”: its use discourages others from also using resistant crop, as they can free-ride off of the efforts of others.

With the default costs of each crop type (where the grower paid *ϕ_T_* = *ϕ_R_* = 0.1), only when tolerant crop is very ineffective (sustaining a high relative loss of yield) do some growers consider resistant crop (Figure 3(A)). Similarly, if resistant crop is very effective (with a low relative susceptibility), there will be a mixed “resistant and unimproved” crop equilibrium (Figure 4). However, due to the free-riding, universal adoption of resistant crop is not achieved, echoing an analogous result in the simpler two strategy model (Murray-Watson & Cunniffe (2022)). Therefore, a mixed equilibrium persists (such as the “unimproved and resistant” equilibrium in Figure 4(A)).

We used the concept of Pareto efficient strategies (Luc (2008)) to optimise the dual objectives of ensuring high profits for growers whilst also minimising the cost to social planners. The positive-feedback induced by the use of tolerant crop means that it will have widespread adoption by growers, so those providing subsidies will have to do so for nearly all growers. This increases the cost to the planner (*τ*), though does increase the profits to the growers (*P*) (Figure 6(E)). However, relatively high profits can also be obtained when resistant crop is subsidised (at least 85% of the maximum theoretical profit, Figure 6(E)). Furthermore, as the adoption of resistant crop is not as widespread as that of tolerant crop, most of the profits can be achieved with less investment from the planner. Therefore, the control strategies on the Pareto front require the planner to provide sufficient subsidies to resistant crop to ensure that it is used whilst never incentivising tolerant crop (Figure 6(A)-(D)). Most optimal subsidisation schemes result in a “unimproved and resistant” crop equilibrium (Figure 6(D)).

By targeting subsidies at resistant crop, the social planner can exploit the freeriding behaviour of growers not growing improved crop and only subsidise a subset of growers. Targeting subsidies to resistant crop can be seen as a way of internalising some of the positive externalities produced by these growers; conversely, not subsidising tolerant crop internalises their negative externalities (Gersovitz (2014)). This strategy was optimal irrespective of the time course over which profits and costs were calculated (Figure 7(F)). However, earlier in the epidemic (between 1-4 seasons), subsidisation schemes that resulted in some fields planted with tolerant crop were possible, but only when the costs to the planner were sufficiently low (Figure 7).

These models ignore the possibility of virus evolution, which is a major threat to the durability of genetic disease control mechanisms (Gallois et al. (2018), Sett et al. (2022)). Tolerant crops, by allowing pathogen replication, do not exert the same selection pressure (Råberg et al. (2009)) and therefore may provide a more durable protection against yield loss (Bingham et al. (2009), Newton (2016), Zhu et al. (2004)). Resistant crop exerts much stronger selection pressures on the pathogen, often leading to “resistance breakdown” where the crop’s resistant traits become ineffective against infection (García-Arenal & McDonald (2003), Parlevliet (2002)). Over the short term, it may be cheaper for the planner to subsidise the use of resistant crop, but when considering the potential evolution of a resistance-breaking pathogen and the associated costs of developing and disseminating new crop types, tolerant crop may be more favourable.

Other simplifying assumptions were made for this study. We did not include spatial or stochastic effects (Cunniffe et al. (2015), Hyatt-Twynam et al. (2017), Cunniffe & Gilligan (2020), Fabre et al. (2021)), both of which can influence disease progression and growers’ decision-making. We also did not differentiate between the nature or quantity of information available to each grower, both of which will depend on the grower’s social and professional network (Milne et al. (2016), Sherman & Gent (2014)) or allow for differing perceptions of risk (our values of responsiveness, *η_b_, b* ∈ {*U,T,R*} were the same across all growers irrespective of control strategy). A grower’s “risk attitude” will be influenced by factors such as their previous experience with disease outbreaks (Garcia-Figuera et al. (2021)) or received knowledge from other growers (Sherman & Gent (2014)) and will impact their control decisions and consequently disease progression (Murray-Watson et al. (2022)). Growers, then, are ultimately “reflexive producers” (Kaup (2008)), and their control decisions must balance each of these information sources with external factors such as market pressures.

Our model demonstrates how decision models of grower behaviour can be combined with other economic concepts, such as Pareto fronts, to find socially-optimal solutions across conflicting objectives. We found that what was optimal for the growers (using tolerant crop) led to worse outcomes for the planner, and the social optimum occurred when a subset of the growers used resistant crop. Though we examine these trade-offs for a crop that is either tolerant or resistant to TYLCV, the model form is sufficiently flexible to allow for other pathosystems or control mechanisms.

## 6 Data availability

Base code is available at: https://github.com/RachelMurray-Watson/Expanding-growers-choice-of-disease-management-options-can-promote-suboptimal-social-outcomes (Murray-Watson (2022)).

## 7 Appendix 1: Calculating the expected profit

To calculate the expected profit, we must first calculate the profits associated with each possible outcome a grower could achieve at the time of harvest (or when a field is rogued). These will depend on the control strategy used by the grower (whether they planted tolerant, resistant, or unimproved crop at the beginning of the season), the infectious status of their field and whether an infectious field was rogued.

The profits for each field type are given as follows:

*P_SU_* = Profit for non-controller with a susceptible field,

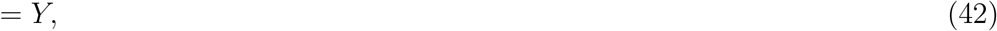

*P_EU_* = Profit for non-controller with latently-infected field,

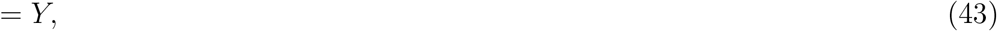

*P_I_H_U_* = Profit for non-controller with an infected field that was not rogued,

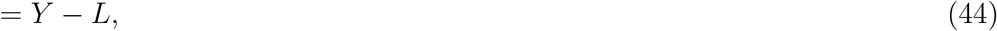

*P_I_R_U_* = Profit for non-controller with an infected field that was rogued,

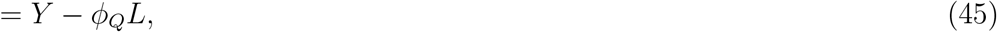

*P_S_*T = Profit for controller using tolerant crop with a susceptible field,

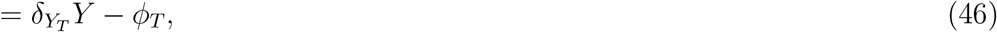

*P_ET_* = Profit for controller using tolerant crop with latently-infected field,

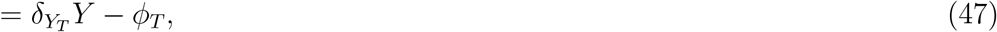

*P_I_H_T_* = Profit for controller using tolerant crop with an infected field that was not rogued,

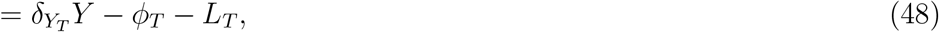

*P_I_R_T_* = Profit for controller using tolerant crop with an infected field and that was rogued,

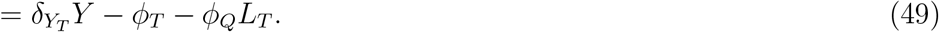

*P_SR_* = Profit for controller using resistant crop with a susceptible field,

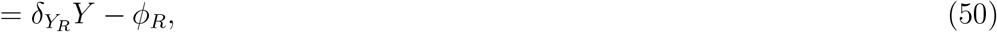

*P_ER_* = Profit for controller using resistant crop with latently-infected field,

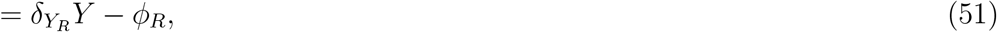

*P_I_H_R_* = Profit for controller using resistant crop with an infected field that was not rogued,

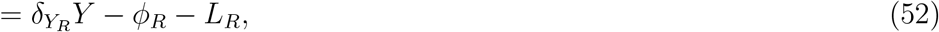

*P_I_R_R_* = Profit for controller using resistant crop with an infected field and that was rogued,

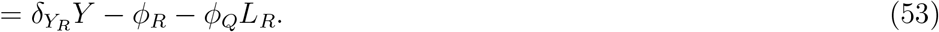

The expected profits for each strategy depend on the probabilities of a grower receiving each of these payoffs.

For each crop variety, the probability of infection (*q_j_*) is given as:

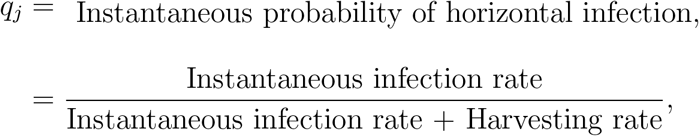

so

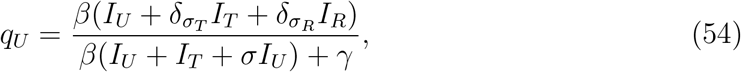

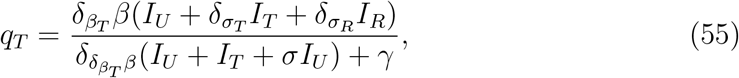

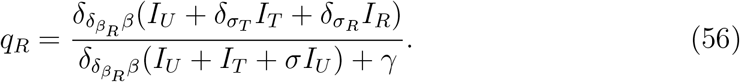

The expected profit, however, will also be dependent on whether a grower’s field is harvested whilst latently infected (*q_Eb_*) or proceeds to become fully infectious (*q_Ib_*).

The probability of being harvested whilst latently infected is given as:

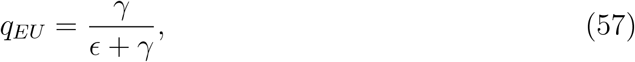

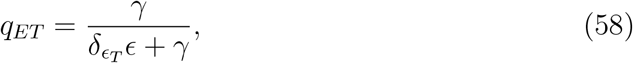

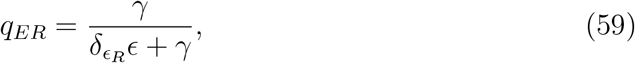

and the probability of becoming fully infectious before harvest is:

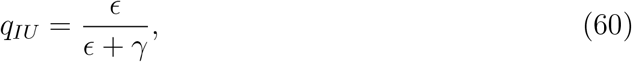

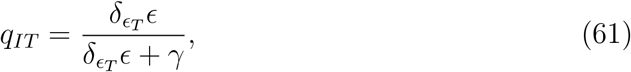

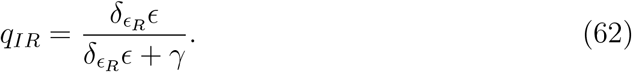

Fully infectious crops can either be harvested or removed via roguing. The probability that harvesting occurs before roguing (*q_I_H_b_*) is given by:

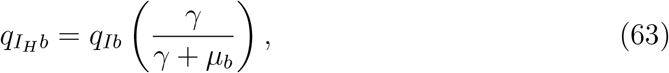

and that it is rogued before harvesting (*q_I_R_b_*) is:

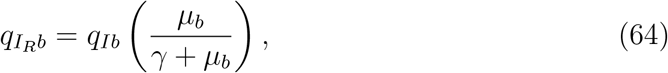

We can then use these probabilities to calculate the expected profit of each strategy:

*P_U_* = Grower’s estimate of the expected profit next season if control is not adopted,

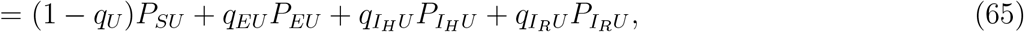

*P_T_* = Grower’s estimate of the expected profit next season if tolerant crop is used,

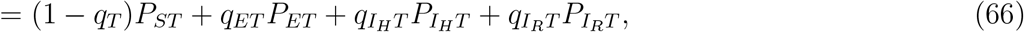

*P_R_* = Grower’s estimate of the expected profit next season if resistant crop is used,

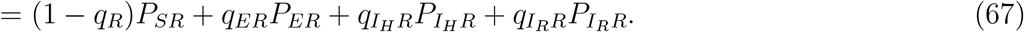

We can simplify these expressions to:

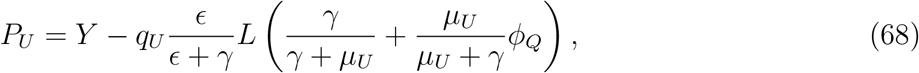

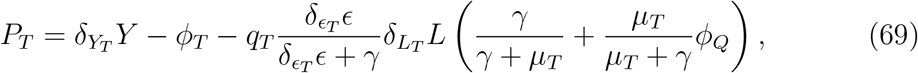

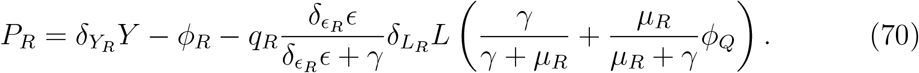

## 8 Appendix 2: Details of switching terms

The switching terms take the general form: a grower with outcome *P_ab_*, is switching into strategy *c* or *d, a* ∈ {*S,E,I_H_,I_R_*} and *b,c,d* ∈ {*U,T, R*},*b* ≠ *c* ≠ *d*. If the profits of both *c* and *d* are exactly equal, the grower will switch into whichever strategy has the higher proportion of current users. If both the expected profits and the proportion of growers using each strategy are the same, then half of the growers considering changing strategy will compare with each alternative expected profit.

The full details of all the switching terms are given below. If *P_T_ < P_R_* or *P_T_* = *P_R_* and *R* > *T* (i.e. there are more using resistant than tolerant crop), then

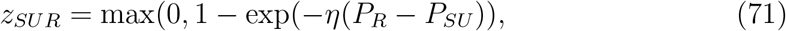

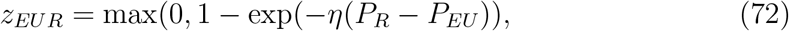

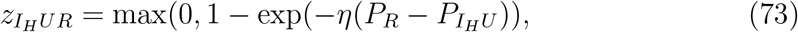

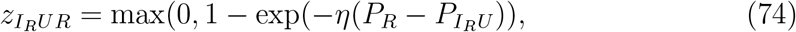

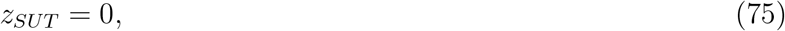

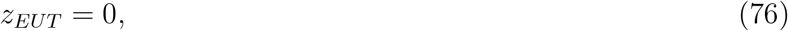

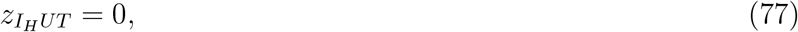

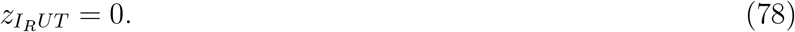

If *P_T_* > *P_R_* or *P_T_* = *P_R_* and *R* < *T*, then

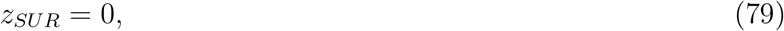

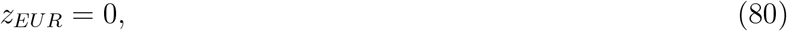

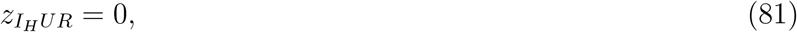

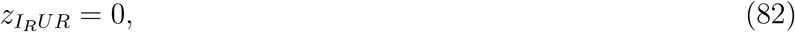

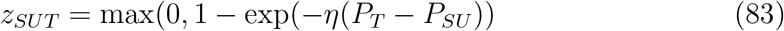

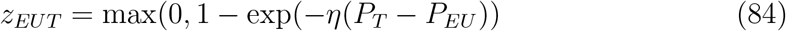

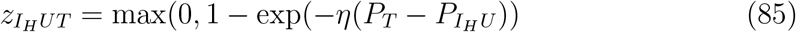

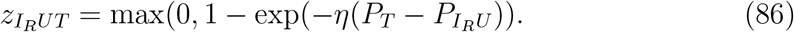

If *P_T_* < *P_U_* or *P_T_* = *P_U_* and *U* > *T*, then

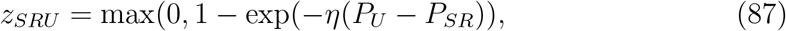

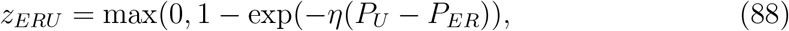

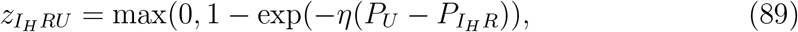

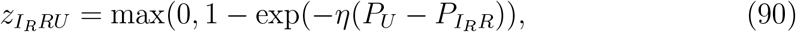

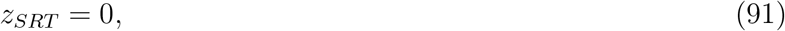

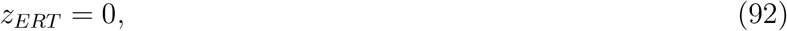

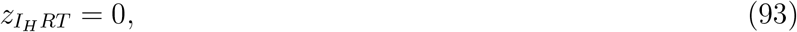

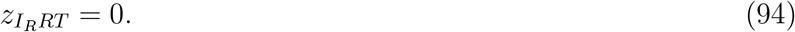

If *P_T_* > *P_U_* or *P_T_* = *P_U_* and *U* <*T*, then

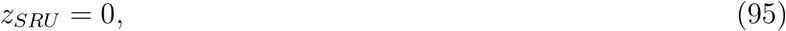

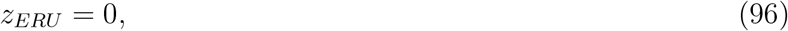

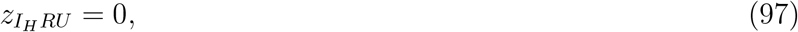

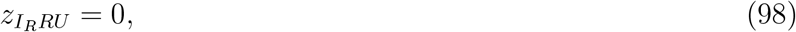

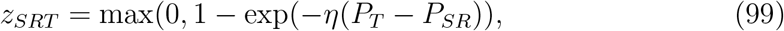

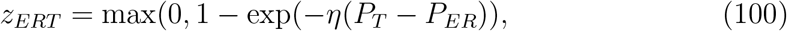

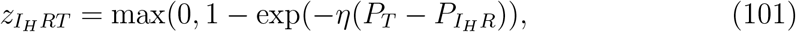

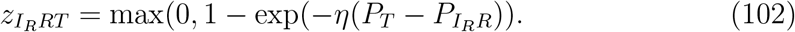

If *P_U_* > *P_R_* or *P_U_* = *P_R_* and *R* < *U*, then

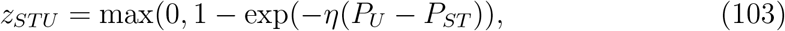

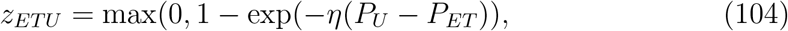

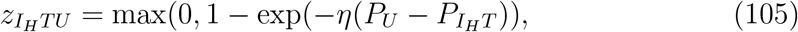

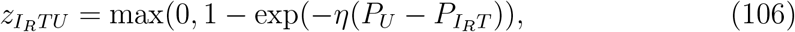

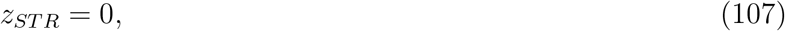

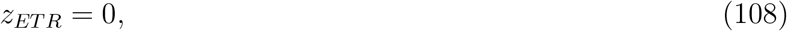

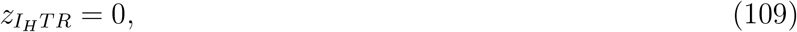

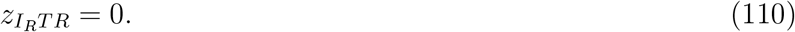

If *P_U_* < *P_R_* or *P_U_* = *P_R_* and *R* > *U*, then

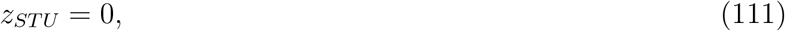

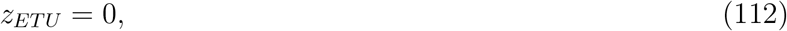

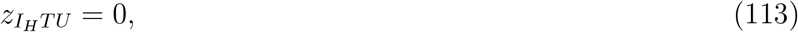

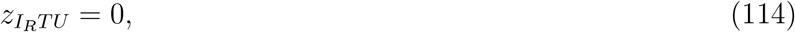

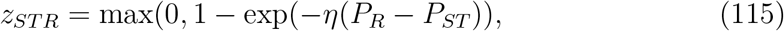

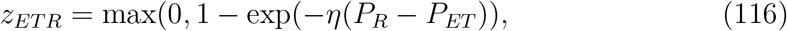

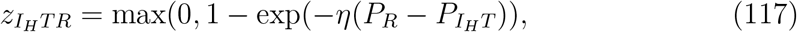

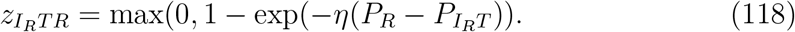

In the case where both the profits and the proportion of growers using the alternative strategy are equal, growers changing strategy will be divided evenly between the two alternative strategies.

To simplify writing the model, we can then say:

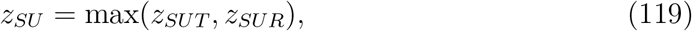

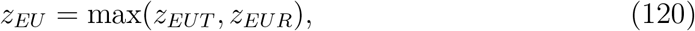

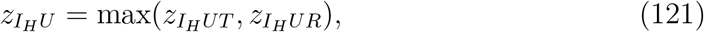

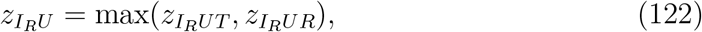

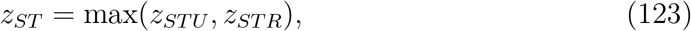

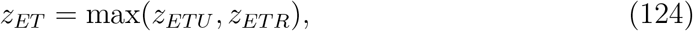

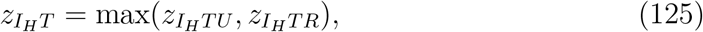

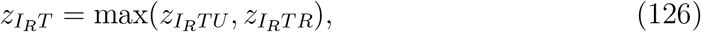

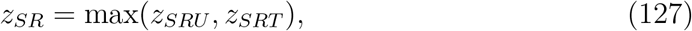

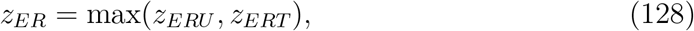

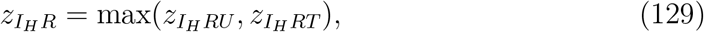

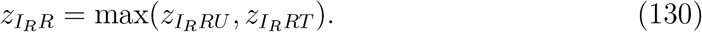

## 9 Appendix 3: Calculating the basic reproduction number for the model

At the disease-free equilibrium, if growers have a choice between strategies they would never chose to pay for control. Thus, in the behavioural model, the disease-free equilibrium is given as: (*S_U_*, *E_U_*, *I_U_*, *S_T_*, *E_T_*, *I_T_*, *S_R_*, *E_R_*, *I_R_*) = (*N*, 0, 0, 0, 0, 0, 0, 0, 0).

For this model, *F* =

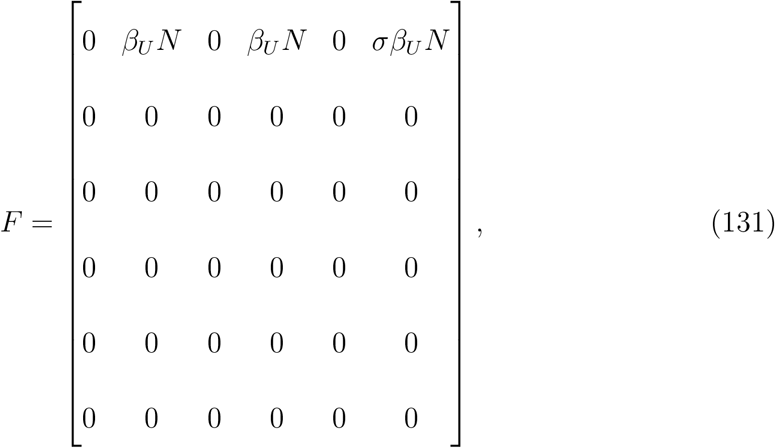

and

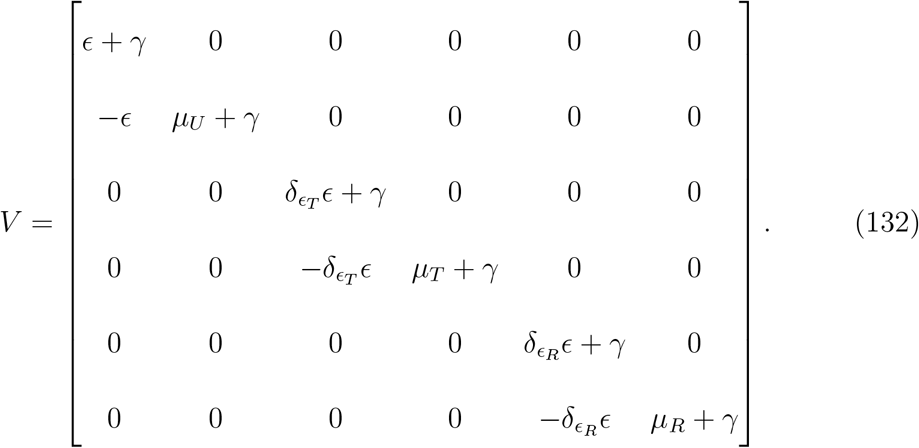

The inverse of *V* is given by:

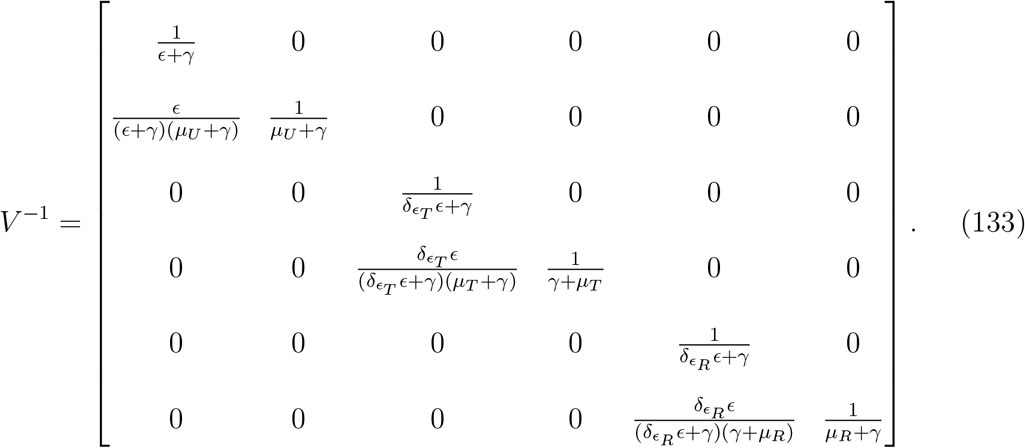

The NGM, *K* = *FV*^−1^, is then given by:

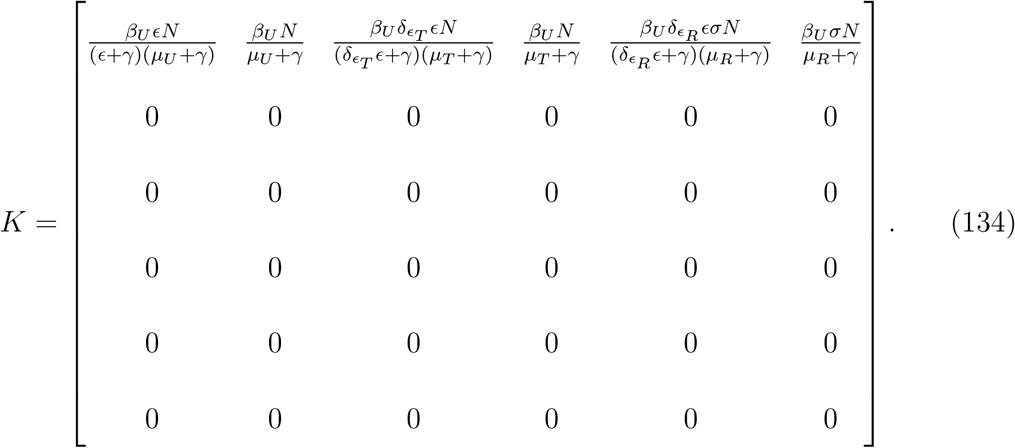

The leading eigenvalue for this matrix is

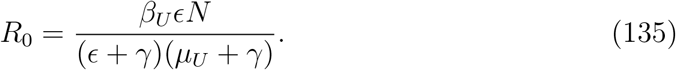

## 10 Appendix 4: Supplementary results

### 10.1 Change in switching points for parameter scans

The underlying bistability of the model means that, for a given set of parameters, different equilibria can be attained depending on the initial conditions. This manifests in our parameter scans over the relative loss due to disease in tolerant crop (*δ_L_T__*) and the relative susceptibility in resistant crop (*δ_β_R__*) (Figures 3 and 4).

In these parameter scans, the initial conditions influence the parameter value at which the system switches from a mixed “tolerant and unimproved crop” equilibrium to a “resistant and unimproved crop” equilibrium in the *δ_L_T__* parameter scan (and vice versa in the *δ_β_R__* scan). When the switch occurs depends strongly on the initial proportion of tolerant crop (Figures 2 and 1, which show the “extreme” parameter values for which the switch can occur). Importantly, this does not change the parameter values where growers of infected tolerant crop should switch strategy, as this occurs within the range of *δ_L_T__* that is unaffected by bistability (Figure 1 (B) and (D)).

**Figure 1:**
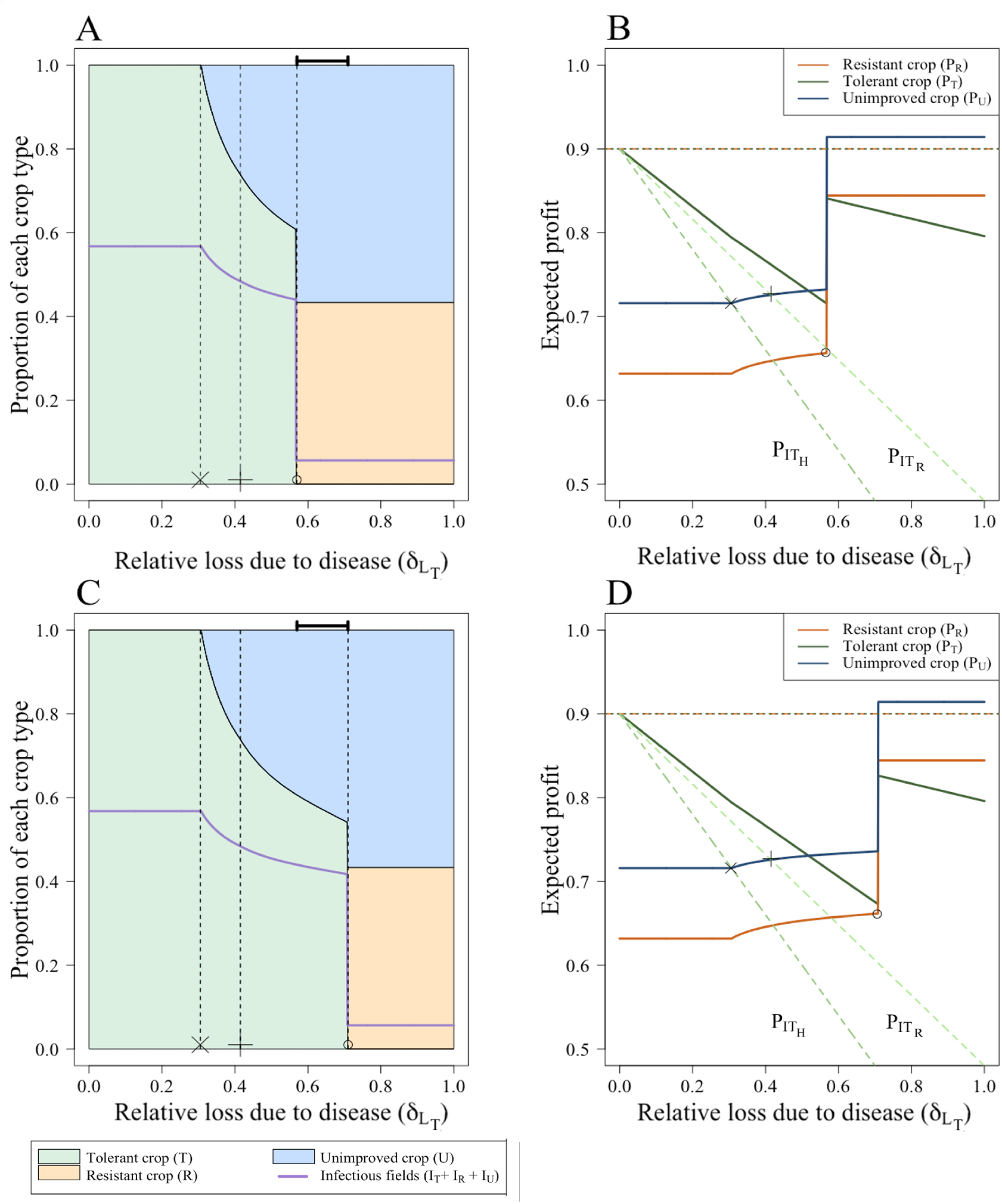
Parameter scans showing range of δ_L_T__ values during which switch in equilibria can occur. (A) + (B) When none of the fields are initially planted with tolerant crop (*T*_0_ = 0), the switch to the “resistant and unimproved” equilibrium occurs at *δ_L_T__* = 0.51 (“O”). (C) + (D) When all of the fields were initially planted with tolerant crop, the switch to the “resistant and unimproved” doesn’t occur until *δ_L_T__* = 0.51 (“O”).

**Figure 2:**
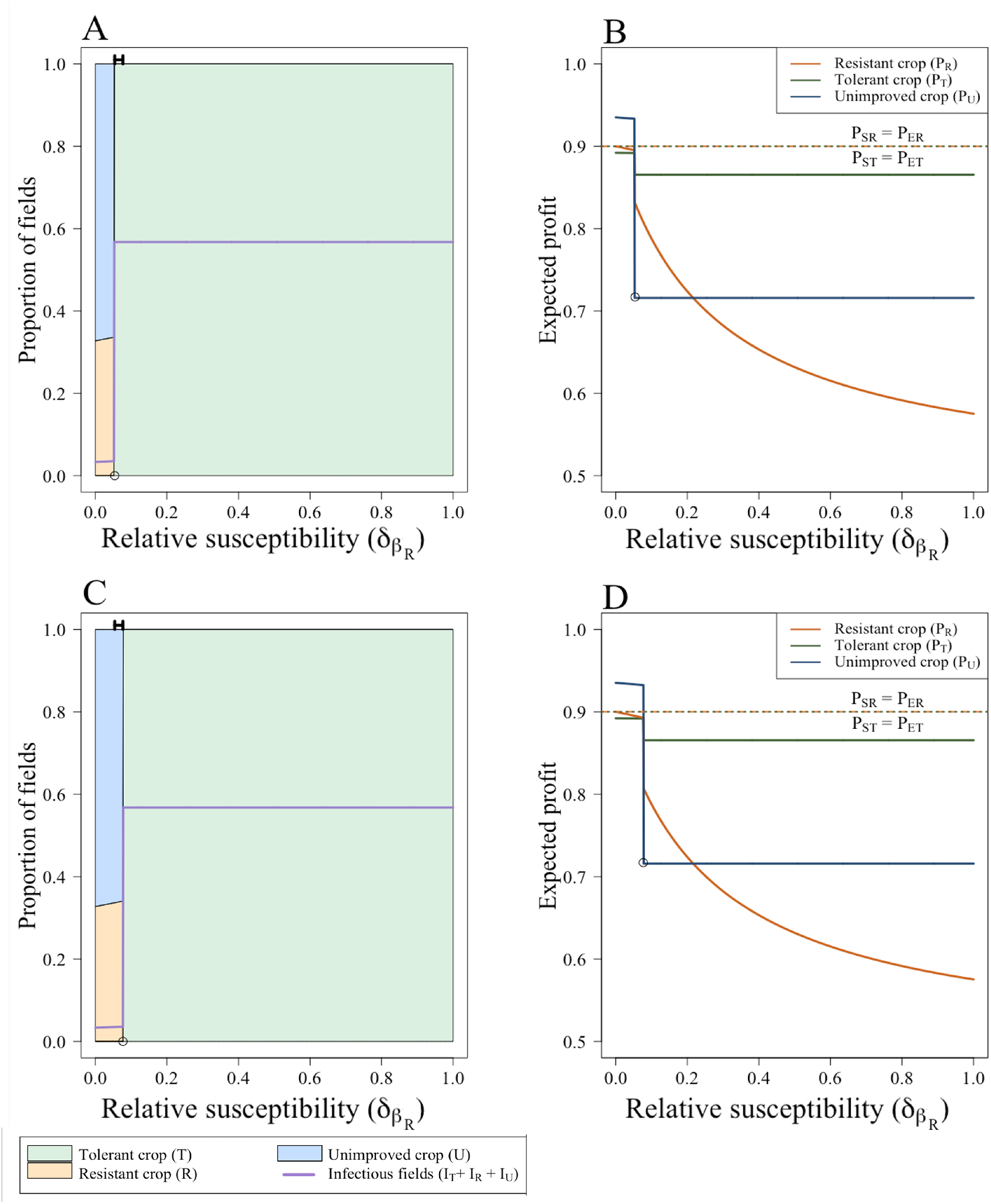
Parameter scans showing range of *δ_β_T__* values during which switch in equilibria can occur. When none of the fields are initially planted with tolerant crop (*T*_0_ = 0), the switch to the “resistant and unimproved crop” equilibrium occurs at a lower value of When all of the fields are initially planted with tolerant crop (*T*_0_ = 0; (A)), the switch to the “resistant and unimproved” equilibrium occurs at lower values of *δ_β_R__* than if *T*_0_ = 1; (B).

### 10.2 Time courses for fairest and least-fair subsidisation schemes

The Pareto front is bounded by two extremes: one where the profit of growers is maximised (*P* = 1), and the other when the cost to the planner is minimised (*τ* = 0). Both of these scenarios have a Gini coefficient of 0.5. To prioritise the growers’ profits, the cost of both resistant and tolerant crop must be low (*ϕ_R_* = 0 and *ϕ_T_* = 0.05). Conversely, to minimise the cost to the planner, both crops must be expensive for the growers (*ϕ_R_* = 0.4 and *ϕ_T_* = 0.375).

The fairest scenario (*G* = 0.009) occurs when *τ* = 0.13 and *P* = 0.93. To achieve this, the cost of tolerant crop (*ϕ_T_*) is 0.12, and the cost of resistant crop (*ϕ_R_*) is 0.105.

Here, we show the dynamics of the model for each scenario. In both the “fairest” scenario and when the profits to the grower are preferred, the subsidisation scheme results in a high proportion using resistant crop and very low levels of disease (Figure 3(A)-(B)). When the costs to the planner are minimised, then no growers use resistant crop, and very few use tolerant crop (which is not subsidised, but still provides a reduced loss in yield if infected).

**Figure 3:**
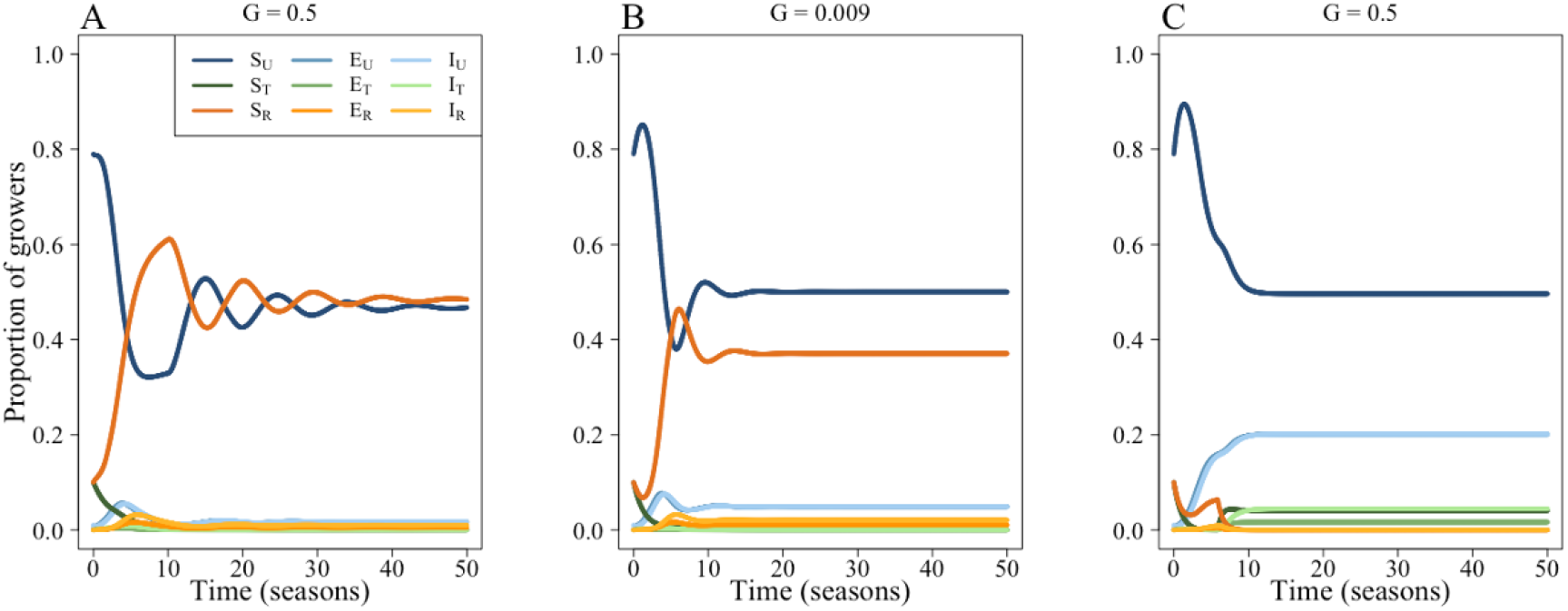
Time courses for the most- and least-fair subsidisation schemes when the tolerant and resistant crop are inefficient. (A) Dynamics that maximise the profits to the grower. Here, ≈ 49.8% of growers use resistant crop. (B) The fairest scenario, where both the profits and costs have relatively equal weighting. In this case, ≈ 40.3% use resistant crop. (C) When minimising the cost to the planner is prioritised, ≈ 10% of growers use tolerant crop.

### 10.3 Effect of change in cost of improved crop on the proportion of tolerant or resistant crops

When the cost of resistant (*ϕ_R_*) and tolerant (*ϕ_T_*) crops are varied, there is only a narrow parameter space where both crop types can coexist (Figure 6(D) in the main text). For the majority of the parameter space, two distinct regions emerge: one where growers use resistant and unimproved crop, and one where growers use tolerant and unimproved crop. The Pareto front lies within the “resistant and unimproved” equilibrium, save for the solution at *ϕ_T_* = 0.375 and *ϕ_R_* = 0.4, where no growers use resistant crop and 10% use tolerant crop.

**Figure 4:**
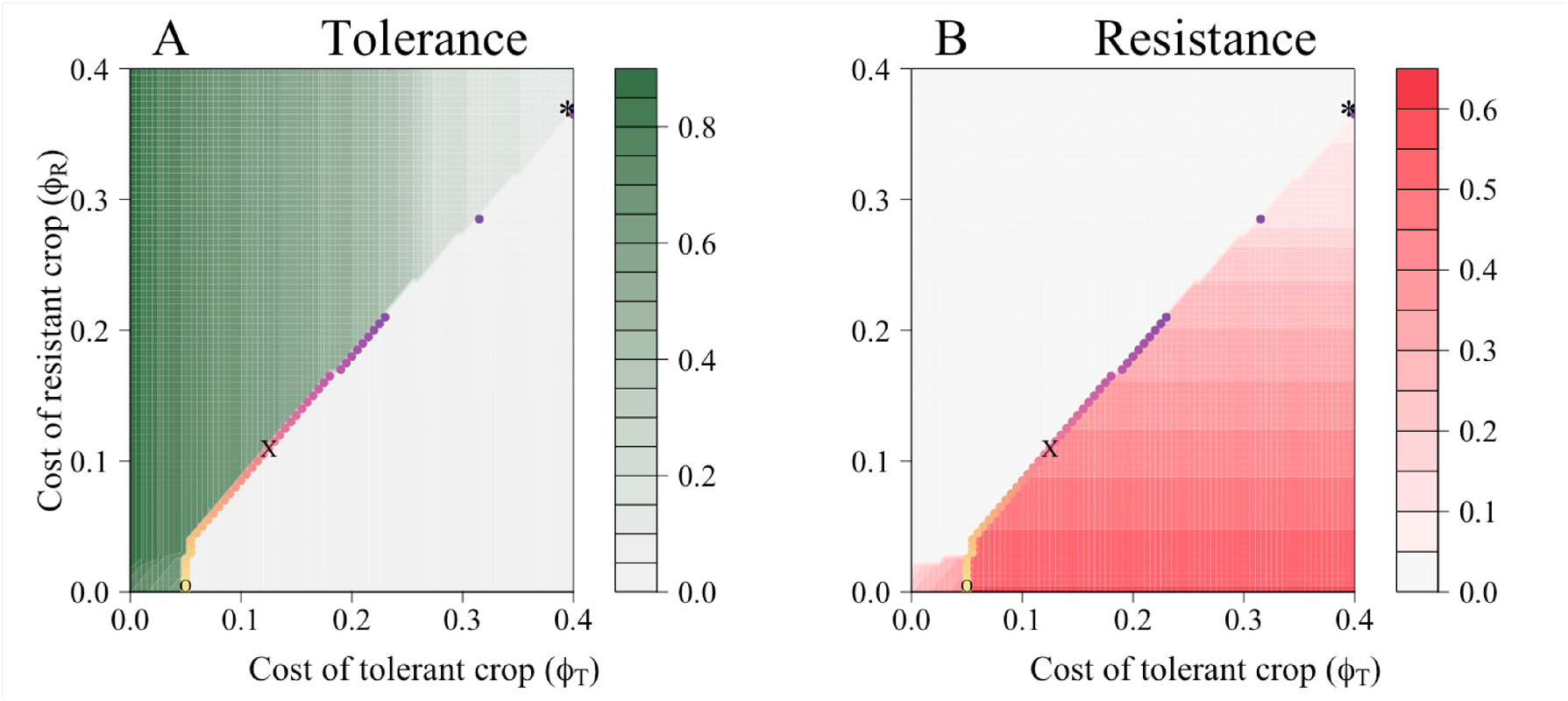
Effect of change in the price of tolerant (*ϕ_T_*) and resistant (*ϕ_R_*) crop on the proportion of growers using improved crop over time. Change in uptake of (A) tolerant crop and (B) resistant crop. Along the Pareto front, growers are always in a “resistant and unimproved” equilibrium, except for the point at *ϕ_T_* = 0.375 and *ϕ_R_* = 0.4, where no growers use resistant crop and 10% of growers use tolerant crop. The fairest scenario is marked with a “X” (Gini coefficient = 0.009, when *τ* = 0.13 and *P* = 0.93). The least fair scenarios (*G* = 0.5) are marked with “O” when the profit is prioritised, and “*” when the costs to the planner are.

